# The polymicrogyria-associated *GPR56* promoter preferentially drives gene expression in developing GABAergic neurons in common marmosets

**DOI:** 10.1101/2020.10.21.348821

**Authors:** Ayako Y Murayama, Ken-ichiro Kuwako, Junko Okahara, Byoung-Il Bae, Misako Okuno, Hiromi Mashiko, Tomomi Shimogori, Christopher A Walsh, Erika Sasaki, Hideyuki Okano

**Author notes:** Current addresses: Department of Neuroscience, University of Connecticut School of Medicine, Farmington, CT, USA. Current addresses: Molecular Analysis for Higher Brain Function, Center for Brain Science, RIKEN, Wako, Japan (Hiromi Mashiko). Ken-ichiro Kuwako and Junko Okahara are joint second authors. Correspondence and requests for materials should be addressed to E.S. and H.O.

## Abstract

GPR56, a member of the adhesion G protein-coupled receptor family, is abundantly expressed in cells of the developing cerebral cortex, including neural progenitor cells and developing neurons. The human *GPR56* gene has multiple presumptive promoters that drive the expression of the GPR56 protein in distinct patterns. Similar to coding mutations of the human *GPR56* gene that may cause GPR56 dysfunction, a 15-bp homozygous deletion in the cis-regulatory element upstream of the noncoding exon 1 of *GPR56* (*e1m*) leads to the cerebral cortex malformation and epilepsy. To clarify the expression profile of the *e1m* promoter-driven GPR56 in primate brain, we generated a transgenic marmoset line in which EGFP is expressed under the control of the human minimal *e1m* promoter. In contrast to the endogenous GPR56 protein, which is highly enriched in the ventricular zone of the cerebral cortex, EGFP is mostly expressed in developing neurons in the transgenic fetal brain. Furthermore, EGFP is predominantly expressed in GABAergic neurons, whereas the total GPR56 protein is evenly expressed in both GABAergic and glutamatergic neurons, suggesting the GABAergic neuron-preferential activity of the minimal *e1m* promoter. These results indicate a possible pathogenic role for GABAergic neuron in the cerebral cortex of patients with *GPR56* mutations.

## INTRODUCTION

G protein-coupled receptor 56 (GPR56) is a member of the adhesion G protein-coupled receptor family^1,2^, and is expressed in multiple tissues including brain, colon, lung, muscle, kidney, pancreas and testis^1^. In the nervous system, GPR56 is highly expressed in neural progenitor cells, and is also expressed in developing neurons^2,3,4,5^. Previous studies have clearly demonstrated the crucial roles of GPR56 in cortical development. In the developing brain, neural stem cells proliferate in the ventricular zone (VZ), and their daughter cells migrate along radial glial fibers toward the pial basement membrane, developing into excitatory glutamatergic neurons. Glutamatergic neurons are generated from deep layer (VI) to upper layer (II) in a birthdate-dependent, inside-out manner. In contrast, GABAergic interneurons originate from the VZ of the ganglionic eminence (GE) and migrate tangentially in multiple streams^6^. Loss of Gpr56 in mice causes many cellular abnormalities in the cerebral cortex, including reduced proliferation of neuronal progenitor cells^7^, structural aberrations in the radial glial endfeet and pial basement membrane, mislocalization of Cajal–Retzius cells, and overmigration of developing neurons^8^. Consequently, *Gpr56*-deficient mice exhibit disorganized cortical lamination and a cobblestone-like malformation^8^. Collagen III α-1, one of the ligands of Gpr56 expressed in the pial basement membrane, is a key molecule involved in Gpr56-mediated neuronal radial migration in the cortex^9^. Upon binding of collagen III, Gpr56 associates with Gα12/13 family of G proteins and activates the RhoA pathway in the radially migrating neurons, leading to properly controlled termination of migration^9^. Gpr56 also plays roles in the proliferation of oligodendrocyte precursor cells and the development and maintenance of peripheral myelin^10,11^.

Consistent with the defects observed in *Gpr56*-deficient mice, multiple *GPR56* coding mutations in human have been found to cause a devastating cortical malformation called bilateral frontoparietal polymicrogyria, as well as frontal lobe-associated dysfunctions, such as epilepsy^2^. The human *GPR56* gene has at least 17 alternative transcription start sites that may drive transcription of mRNAs with different noncoding first exons; these mRNAs encode identical GPR56 protein with distinct expression profiles in the brain^7^. A 15-bp deletion within a cis-regulatory element upstream of the transcriptional start site of noncoding exon 1m (e1m) of *GPR56* has recently been identified in individuals with polymicrogyria restricted to the regions around the Sylvian fissure^7^. All patients with this 15-bp deletion suffer from epilepsy from a young age, as well as from intellectual and language difficulties, but without evident motor disabilities. Epilepsy in patients with the GPR56 mutation is often drug-resistant. Recently, Vigabatrin, γ-vinyl-GABA (a structural analogue of GABA), has been reported to be an effective epilepsy treatment for GPR56-mutated patients^12^. The mechanism of action of Vigabatrin is known to be different from that of GABA_A_R activators such as benzodiazepines and barbiturates, but other details are unknown. The deleted 15-bp sequence is conserved among placental mammals, suggesting a major regulatory element for *GPR56* expression. Given that the patients with the 15-bp deletion in the cis-regulatory element of e1m promoter show a milder and more restricted malformation of the brain compared to patients with coding mutations (which appear to be null mutations), a detailed expression profile of e1m promoter in the cerebral cortex may provide an important insight into the cell types that are responsible for cortical malformation and related symptoms. However, the cell type profile of e1m promoter-driven GPR56 has yet to be characterized.

The common marmoset (*Callithrix jacchus*) has gained prominence as an experimental animal model in the neuroscience field, due to their human-like behaviors and brain structure^13,14,15^. Socially, marmosets form a unit of a close-knit family based on a pair of one male and one female, which is not observed in other experimental model primates, and communicates each other closely through vocalization and eye-contact^16^. In addition, marmoset brain shares structural similarity with human brain, such as the Sylvian fissure and calcarine sulcus, although marmoset is a near-lissencephalic primate^17,18,19,20^. Furthermore, genetics approaches are available thanks to the generation and successful germline transmission of transgenic marmoset^21,22^. This technique enables significant advancements potentially useful in the study of neurological and psychiatric disorders^13,23^.

In the present study, we generated a transgenic marmoset expressing a 0.3-kbp human *GPR56* e1m promoter-driven EGFP (enhanced green fluorescence protein) and provide evidence for preferential activity of the e1m promoter in GABAergic neurons in the developing cerebral cortex, whereas the total GPR56 protein is evenly expressed in GABAergic and glutamatergic neurons as well as progenitor cells. These findings imply a possible role for GABAergic neurons in *GPR56* mutation-associated epilepsy.

## RESULTS

### Production of transgenic marmosets expressing EGFP under the control of human *GPR56 e1m* promoter

To investigate how mutations or deletions within a cis-regulatory element upstream of the e1m of human *GPR56* leads to symptoms in patients, we sought to characterize the cells expressing GPR56 under the control of this cis-regulatory element in marmoset. For this purpose, we generated transgenic marmosets expressing enhanced green fluorescence protein (EGFP) driven by the cis-regulatory element. A previous study reported that 0.3 kb sequence upstream of the human e1m acts as a minimum promoter of the *GPR56* during embryonic stage^7^. The relevant human and marmoset sequences share 92.4% identity, while human and mouse sequences share 62.1% identity in this region (Fig. S1). Regarding the 15-bp element, human and marmoset differ by two bases, while human and mouse differ by one base (Fig. S1). We constructed self-inactivating lentiviral vector harboring this 0.3 kb sequence followed by EGFP coding sequence (referred to hereafter as *0.3k hGPR56 e1m*-EGFP vector) (Fig. 1A). Marmoset zygotes were obtained by *in vitro* fertilization (IVF). Forty-one zygotes were injected with high titer lentiviral vector carrying the *0.3k hGPR56 e1m*-EGFP, of which 27 (65.9%) developed beyond the 4-cell stage (Table 1). Because the *hGPR56* e1m promoter was not active in marmoset preimplantation embryos, the transgene-positive embryos could not be selected by EGFP fluorescence (Fig. S2A), all 27 embryos developed beyond the 4-cell stage were transplanted into 11 recipient females at various developmental stages (Table 1). Five recipient females became pregnant (45.5%) and ultimately two newborns (7.4%. two singletons; one female (I651TgF)) and one male (I757TgM) were delivered naturally at full term (Fig. 1B).

**Table 1.**
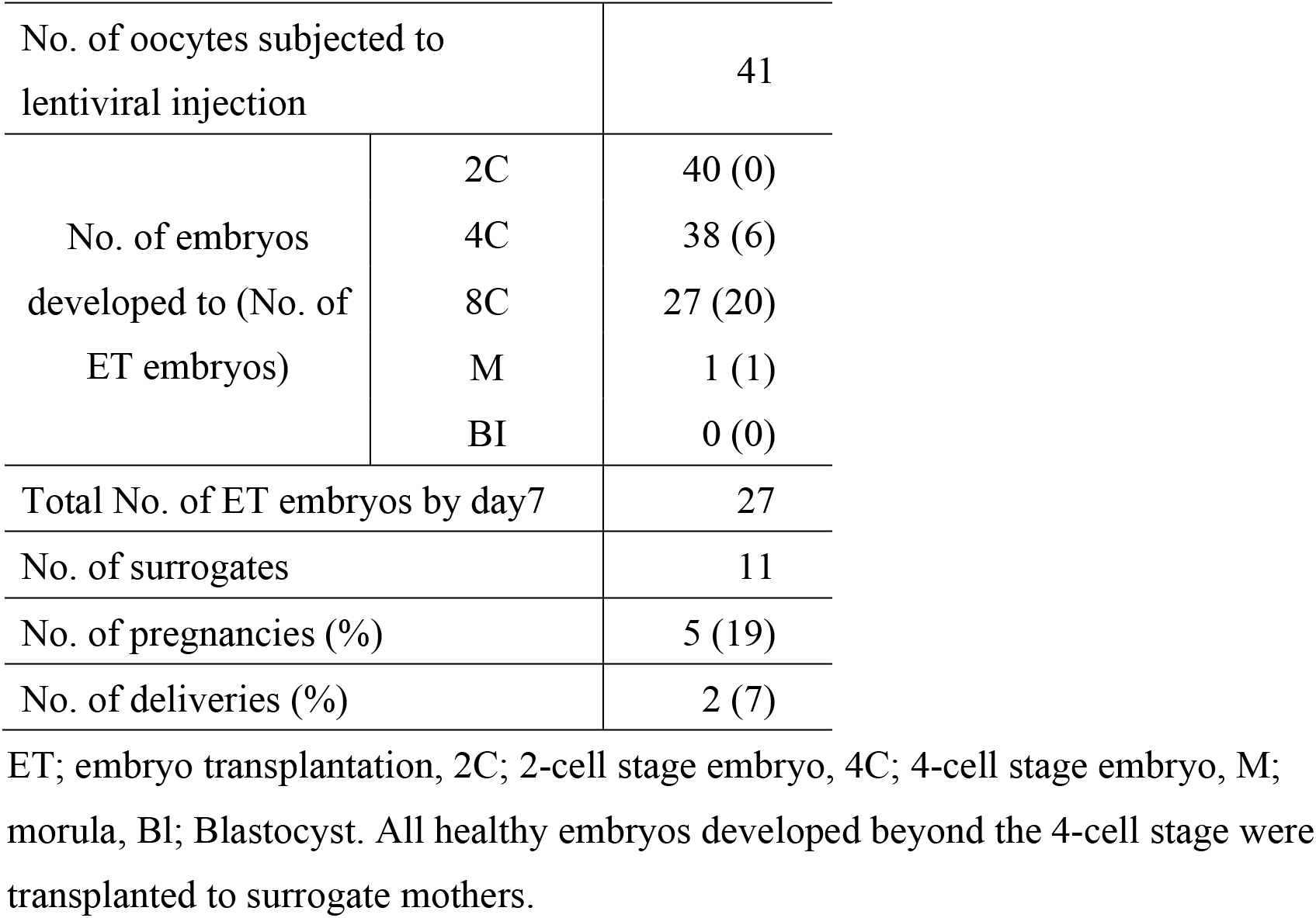
Summary of production of founder (F0) transgenic marmosets.

|  |  |  |
| --- | --- | --- |
| No. of oocytes subjected to lentiviral injection |  | 41 |
| No. of embryos developed to (No. of ET embryos) | 2C | 40 (0) |
|  | 4C | 38 (6) |
|  | 8C | 27 (20) |
|  | M | 1 (1) |
|  | BI | 0 (0) |
| Total No. of ET embryos by day7 |  | 27 |
| No. of surrogates |  | 11 |
| No. of pregnancies (%) |  | 5 (19) |
| No. of deliveries (%) |  | 2 (7) |

ET; embryo transplantation, 2C; 2-cell stage embryo, 4C; 4-cell stage embryo, M; morula, BI; Blastocyst. All healthy embryos developed beyond the 4-cell stage were transplanted to surrogate mothers.

**Figure 1.**
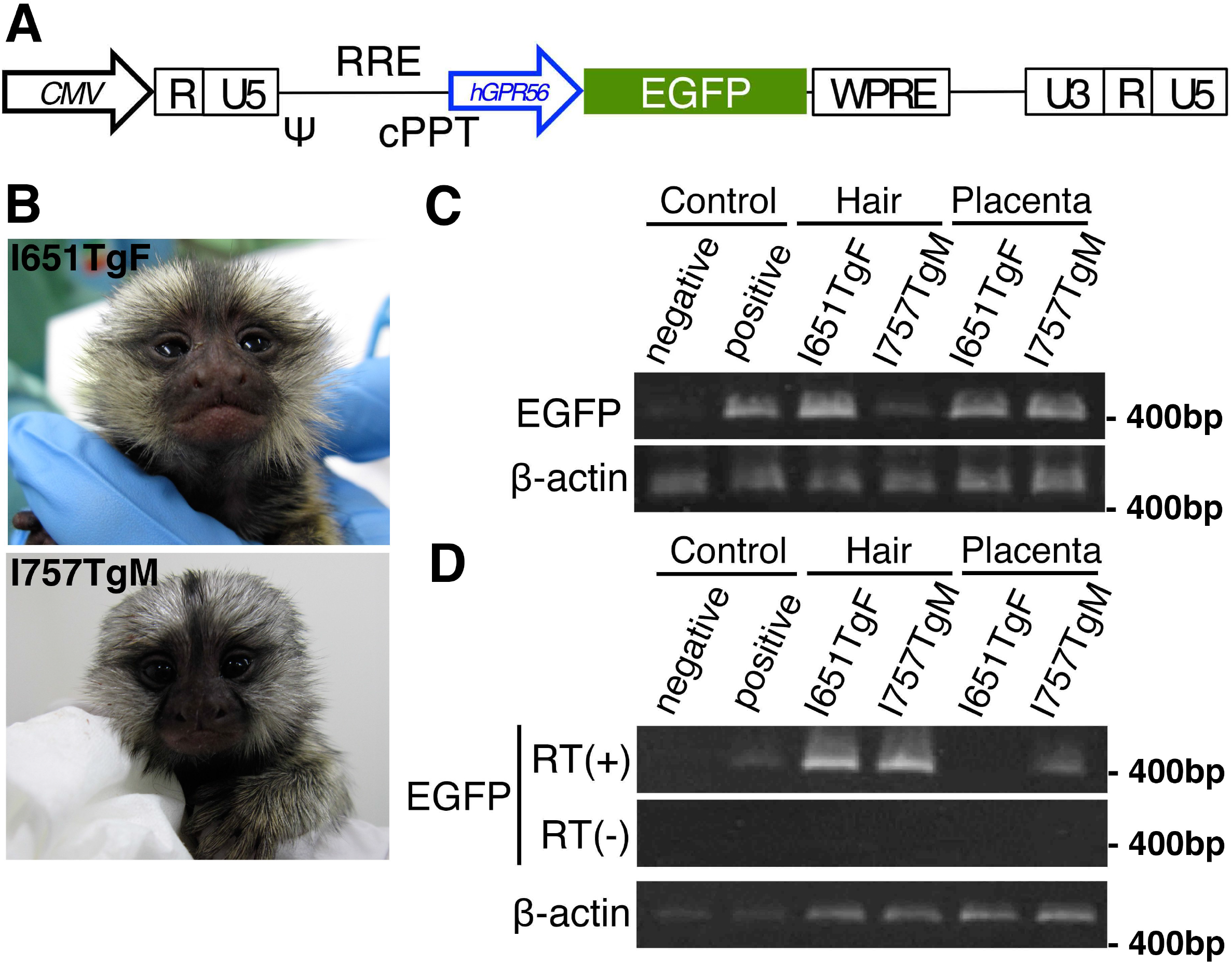
Generation of *0.3k hGPR56 e1m*-EGFP transgenic marmosets. **(A)** Schematic diagram of the lentiviral vector. CMV; Cytomegalovirus promoter, ψ; packaging signal, RRE; rev responsive element, cPPT; central polypurine tract, WPRE:Woodchuck hepatitis virus Posttranscriptional Regulatory Element, *hGPR56*; 0.3-kbp human *GPR56 e1m* cis-element (minimal promoter). **(B)** The transgenic founder infants; I651TgF (female) and I757TgM (male). **(C)** Detection of genomic integration of transgene by genomic PCR. EGFP encoding sequence was amplified using template genomic DNA purified from hair cells of or placentas delivered with infants. Beta-actin amplifications were used as control. (D) Detection of transgene expression by RT-PCR. EGFP encoding sequence was amplified from cDNA templates prepared from RNAs purified from hair cells or placentas delivered with them with (RT+) or without (RT−) reverse transcriptase. Beta-actin amplifications were used as control.

### Transgene integration in the genome

We first examined the genomic integration of the transgene in the infant marmosets. The presence of transgene was tested by PCR using genomic DNA purified from placenta delivered with I651TgF and I757TgM and hair roots of I651TgF and I757TgM. Integrated transgene was detected in all samples (Fig. 1C). Furthermore, EGFP transcripts were detected in hair cells of both infants and in placenta delivered with I651TgF, but not in placenta delivered with I757TgM (Fig. 1D). To identify the chromosomal transgene integration sites, fluorescence *in situ* hybridization (FISH) was performed. There were 27 transgene integration sites on chromosomes 1, 2, 3, 4, 9, 10, 11, 13, 15, 17, 18, 21 and 22 in the peripheral blood cells of I651TgF, while I757TgM had 13 transgene integration sites on chromosomes 1, 2, 4, 6, 7, 11, 13, 15, 16 and 17 (Fig. S2B). These results indicate that I757TgM and I651TgF are transgenic marmosets harboring functional *0.3k hGPR56 e1m*-EGFP transgene in their genome.

### Germline transmission of the transgene

After sexual maturation, the female animal (I651TgF) was mated with a non-transgenic male and 19 embryos were obtained by uterine flushing. Sixteen of these were transferred into eight surrogate mothers(Table 2), and then three F1 fetuses at E95, two at E113, and one at E126 were obtained by caesarean section. EGFP fluorescent signals were detected in all fetuses (Fig. 3A, Fig. 4A, Fig. S3A, B), implying that transgenes were transmitted through oocytes of I651TgF. As for I757TgM, the presence of transgene was detected in the sperm sample by genomic PCR (Fig. S3C). On the other hand, we paired the male transgenic marmoset, I757TgM, with wild-type female and performed uterine flushing to obtain embryos. Although the sperm concentration and motility were normal in the semen of I757TgM, no zygotes were obtained from the I757TgM line for unknown reasons (Table 2).

**Table 2.** Summary of embryo production from F1 transgenic marmosets.

| Parents | I651TgF x WT | WT x I757TgM |
| --- | --- | --- |
| No. of procedures | 15 | 13 |
| No. of collected embryos | 19 | 0 |
| No. of transplanted embryos | 16 | 0 |
| No. of surrogates | 8 | 0 |
| No. of fetuses obtained by caesarean section | 6 | 0 |

**Figure 2.**
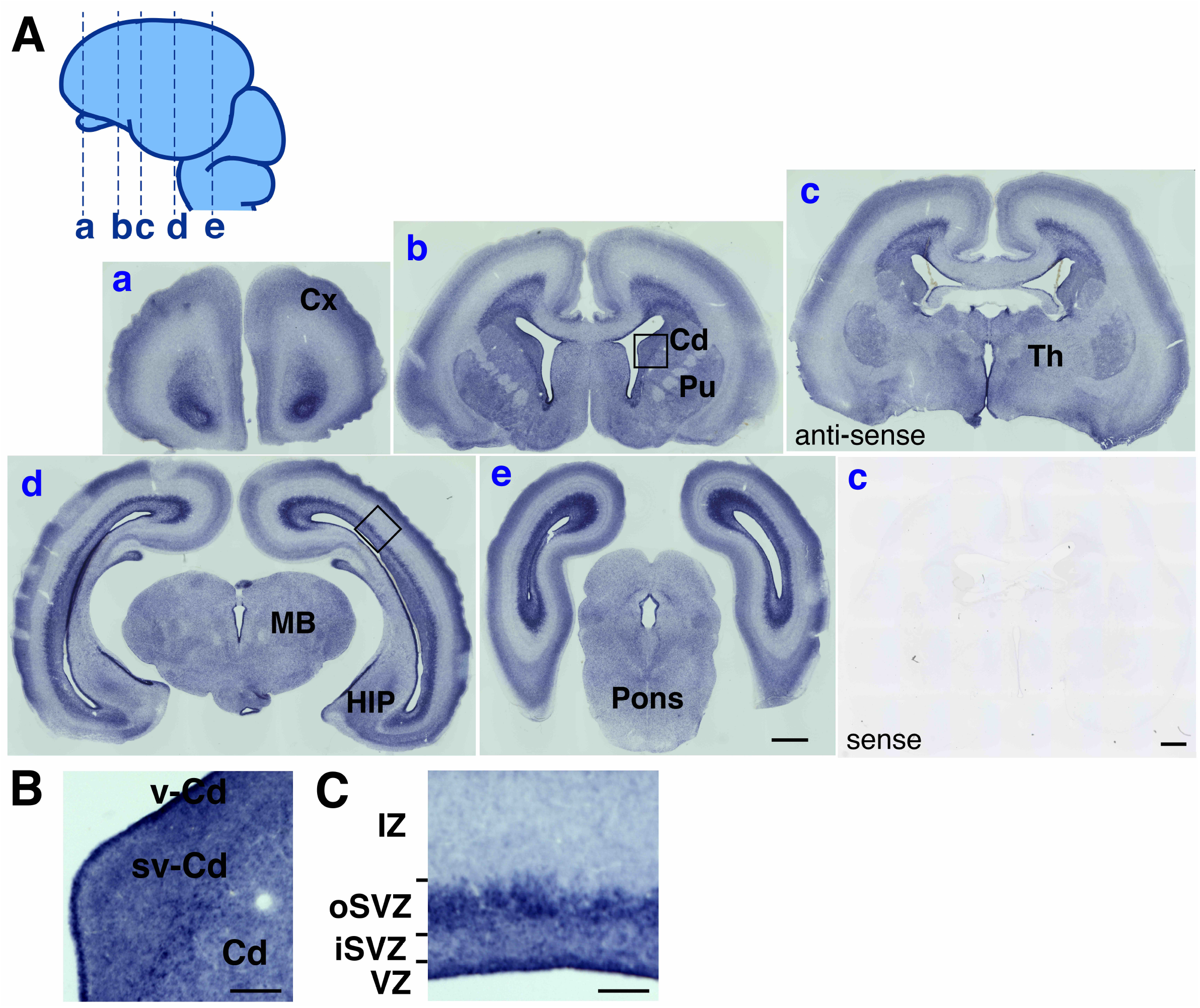
Expression of *GPR56* mRNA in the brain of wild type marmoset embryo. **(A)** Coronal sections of transgenic marmoset brain at the fourteenth embryonic week (EW) were hybridized with anti-sense (a-e) or sense probe (c) for *GPR56* mRNA. Cerebral cortex (Cx), caudate nucleus (Cd), ventricular zone of the caudate (vCd), subventricular zone of the caudate (svCd), thalamus (Th), hippocampus (HIP), pons, midbrain (MB), ventricular zone (VZ), inter subventricular zone (iSVZ), outer subventricular zone (oSVZ), and intermediate zone (IZ) are indicated. Scale bars =1 mm. **(B)** Enlarged image of the area marked by a square in panel b in (A). **(C)** Enlarged image of the area marked by a square in panel d in (A). Scale bar in B and C = 200 μm.

**Figure 3.**
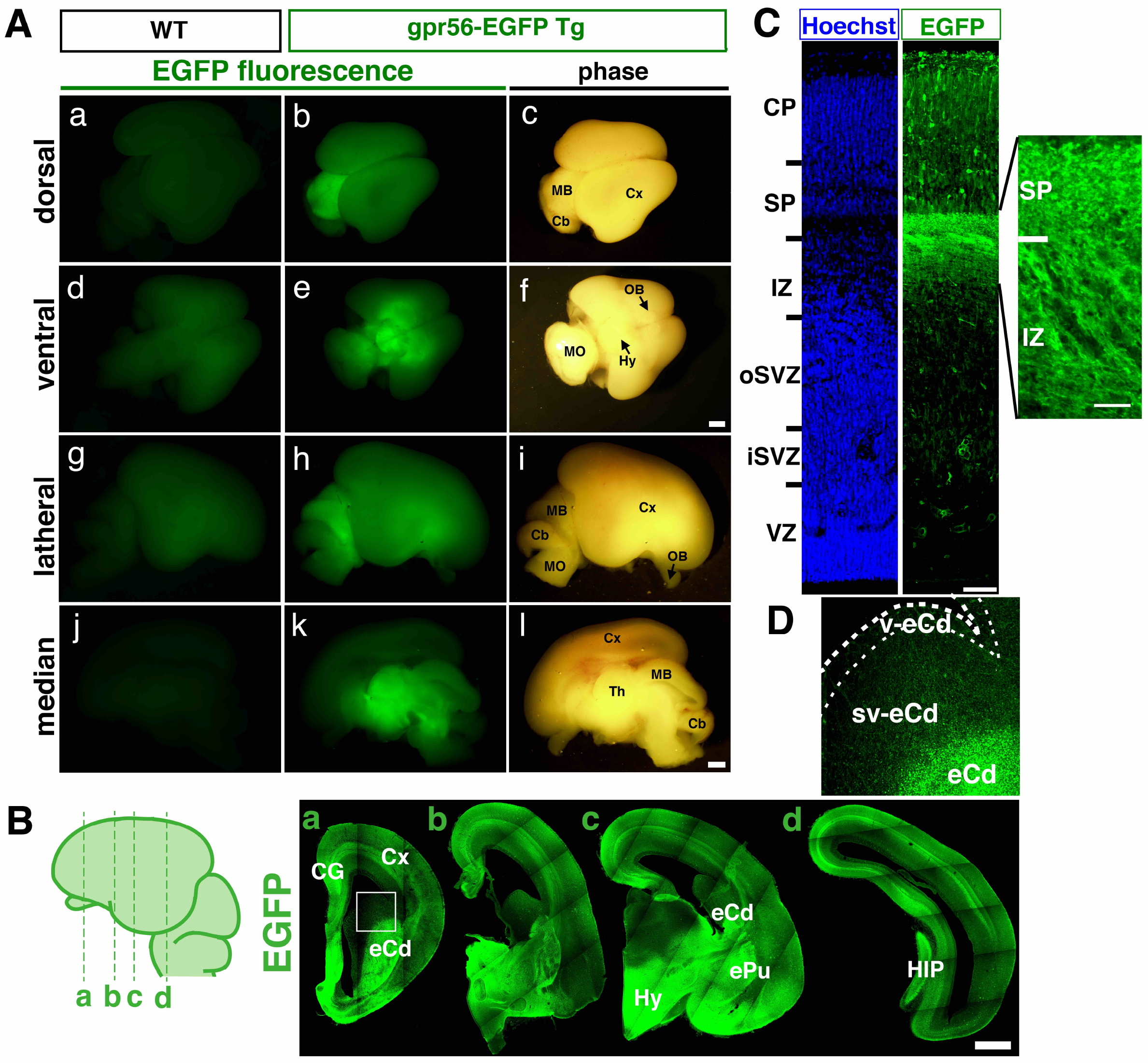
Expression of *hGPR56 e1m*-driven EGFP in transgenic marmoset embryo. **(A)** EGFP signals in the Dorsal view (**a-c)**, ventral view (**d-f)**, lateral view **(g-i)**, and median view **(j-l)** of the whole brain of wild type (left panels) and transgenic marmoset (middle and right panels) at E95. Thalamus (Th), hypothalamus (Hy), midbrain (MB), cerebellum (Cb), medulla oblongata (MO), cerebral cortex (Cx), and olfactory bulb (OB) are shown. **(B)** Immunofluorescent staining of coronal sections from the anterior to posterior (d) of the transgenic marmoset brain (E95) for EGFP. Lines on the schema indicate the position of each section. Cingulate gyrus (CG), early caudate (Cd), early putamen (Pu), hippocampus (HIP). Scale bars in A and B = 1 mm. **(C)** Coronal section of cerebral cortex of the transgenic marmoset brain (E95) stained for DNA (Hoechst) and EGFP. Cortical plate (CP), subplate (SP), IZ, oSVZ, iSVZ, and VZ are indicated. Scale bar = 200 μm. **(D)** Enlarged image of the area marked by a square in the panel of (B), with indication for Cd, ventricular zone of Cd (vCd), and subventricular zone of Cd (svCd).

**Figure 4.**
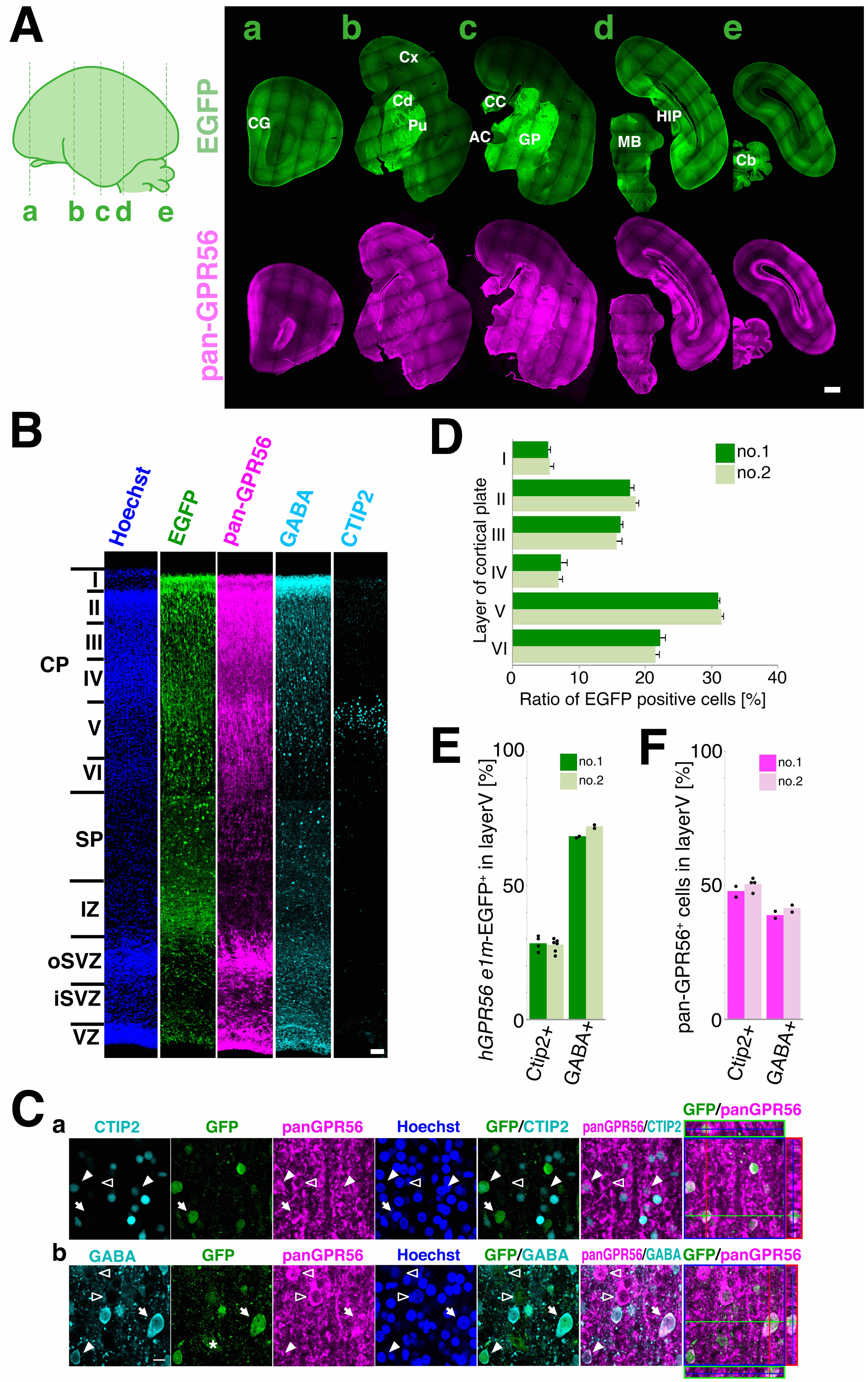
*hGPR56 e1m*-EGFP expression in the cerebral cortex. **(A)** Immunofluorescent staining of coronal sections from the anterior (a) to posterior (e) of the transgenic marmoset brain at E113 for EGFP (green) and pan-GPR56 (magenta). Lines on the schema indicate the position of each section. Scale bar = 1mm. **(B)** Coronal section of cerebral cortex of the transgenic marmoset brain at E113 stained for Hoechst (Blue), *hGPR56 e1m*-EGFP (green), and pan-GPR56 (magenta), CTIP2 or GABA (cyan). The section stained for CTIP2 is the one 50 μm anterior to the other sections. Cortical plate (CP) consisting of layers I to VI, SP, IZ, oSVZ, iSVZ, and VZ are indicated. Scale bar = 200μm. **(C)** Distribution of GPR56 (magenta), *hGPR56 e1m*-EGFP (green), and CTIP2 or GABAs (cyan) positive cells in layer V of the cortical plate. Representative *hGPR56 e1m*-EGFP positive cells with (filled arrowheads) or without (open arrowheads) expression of CTIP2 or GABAs are marked. The panels furthest to the right display orthogonal views of a *hGPR56 e1m*-EGFP positive cell marked by the arrow. Orthogonal views of other *hGPR56 e1m*-EGFP positive cells are shown in Figure S5A. We used ZEN 2009 software (version: 6.0.0.303, Carl Zeiss, Oberkochen, Germany) to construct orthogonal views. Scale bar = 50 μm. **(D)** Ratio of *hGPR56 e1m*-EGFP positive cells in each CP layer of transgenic marmoset no.1 and no.2. **(E)** Ratio of CTIP2^+^ or GABA^+^ cells among the *hGPR56 e1m*-driven EGFP^+^ cells in layer V of transgenic marmosets no.1 and no.2. **(F)** Ratio of CTIP2^+^ or GABA^+^ cells among the pan-GPR56^+^ cells in layer V of transgenic marmosets no.1 and no.2.

### Expression pattern of endogenous GPR56 and *0.3k hGPR56 e1m*-driven EGFP in the embryonic brain

GPR56 is expressed in various tissues including brain in human and mouse^7^. We confirmed broad expression of GPR56 in embryonic marmoset brain by *in situ* hybridization. At the 10th, 12th and 14th embryonic week (EW), strong GPR56 signals were detected in VZ, mainly consisting of neural stem cells. At the 14th EW, GPR56 was also strongly expressed in outer subventricular zone (oSVZ), where the highly proliferative progenitor cells called basal radial glia reside (Fig. 2 and Fig. S4) ^24,25,26^. These results suggest abundant *GPR56* mRNA expression in immature neural cells.

In the F1 fetus brain at E95, *0.3k hGPR56 e1m*-driven EGFP protein expression was observed in restricted areas, such as thalamus, hypothalamus, midbrain, cerebral cortex (Fig. 3A). In order to determine the EGFP expression pattern in more detail, we prepared brain slices from E95 transgenic marmoset embryos in the coronal plane and stained them with anti EGFP antibody. The signals were mainly detected in cerebral cortex, cingulum, early caudate nucleus, early putamen, hippocampus, and hypothalamus (Fig. 3B). In the cerebral cortex, EGFP signals were found in a subpopulation of cells in the cortical plate, as well as in most of the nerve fibers in the subplate and intermediate zone (Fig. 3C). Interestingly, only a few EGFP-positive cells were sparsely distributed in inner subventricular zone (iSVZ) and oSVZ in the cerebral cortex and subventricular zone of the early caudate nucleus and no EGFP-positive cell was found in VZ in the cerebral cortex or VZ of the early caudate nucleus (Fig. 3D), despite the high expression of endogenous *GPR56* mRNA in these zones (Fig 2). Taken together, these results suggest that *0.3k hGPR56 e1m* contains a cis-element that promotes predominant expression of GPR56 in the GE and in a subset of developing neurons in the cortical plate of the marmoset fetus brain.

### EGFP expressed predominantly in GABAergic neurons

To determine the cell types that express *0.3k hGPR56 e1m*-driven EGFP in developing cortex, we performed immunohistochemistry with the cerebral sections at E113, as all layers in CP became distinguishable at this developmental stage. We first confirmed the reactivity of anti-pan-GPR56 antibody with marmoset GPR56 by immunofluorescence using COS cells expressing marmoset GPR56 (Fig. S5). Then, we stained coronal sections with anti-EGFP and anti-pan-GPR56 antibodies and overviewed the sections from rostral to caudal. Similar to at stage E95, EGFP signals were mainly detected in cerebral cortex, cingulum, caudate nucleus, putamen, globus pallidus, hippocampus, hypothalamus and cerebellum, while the pan-GPR56 signals were also strongly observed in VZ and oSVZ (Fig. 4A and 4B). In the cortical plate, 89.6%±1.1(mean±SEM) of the EGFP positive cells (1278 cells in the 10 sections derived from 2 embryos) were positive for pan-GPR56 staining. Next, we examined the distribution of EGFP-positive cells in each cortical layer of two transgenic marmosets, no.1 and no.2. EGFP-positive cells were detected in all layers at various ratios, and were most abundant in the layer V (Fig. 4D and Table S1). We evaluated the ratio of excitatory and inhibitory neurons among EGFP-positive cells by staining GABA and CTIP2, markers for GABAergic inhibitory neurons and for glutamatergic excitatory neurons in deep layer (layer V and VI) ^27^, respectively (Fig. 4B, 4C, 4E, 4F, Fig. S6 and Table S2). In layer V of E113 marmosets, on average, 70.0% and 28.1% of *0.3k hGPR56 e1m*-driven EGFP cells were GABA-positive and CTIP2-positive, respectively (Fig. 4E and Table S2). In contrast, among pan-GPR56 positive cells in layer V of marmosets at E113, 40.3% and 49.1% on average were GABA-positive and CTIP2-positive, respectively (Fig. 4F and Table S2). These results suggest that the *hGPR56 e1m* drives protein expression preferentially in GABAergic neurons rather than glutamatergic neurons in layer V. In other layers of cortical plate, EGFP-positive cells contained a higher percentage of GABAergic neurons than pan-GPR56 positive cells (Fig. S7, Table S3 and S4). At E89, an earlier stage, pan-GPR56 positive cells included almost all Nkx2.1 positive progenitor cells in presumptive MGE (medial ganglionic eminence), from which GABAergic neurons originate (Fig. S8A)^28^. At E95, EGFP positive migrating neurons were observed beneath and within the intermediate zone (Fig. S8B). These findings are consistent with the fact that hGPR56 e1m drives protein expression preferentially in GABAergic neurons. Furthermore, we examined the subtype of the EGFP-positive GABAergic interneurons by analyzing the expression of the principal subtype markers, such as parvalbumin (PV), somatostatin (SST), or calretinin (CR) (Fig. 5A). Of the EGFP-expressing cells in layer V, 50.0% (n=152) on average was PV-positive, 20.0% (n=236) was SST-positive, and 21.9% (n=321) was CR-positive neuron (Fig. 5B and Table S5). While at E113, PV-positive, SST-positive, and CR-positive neurons were found at 34.7% (n=471), 9.2% (n=218), and 10.6% (n=405) of whole cells in layer V, respectively (Fig. 5C and Table S6). These results suggest that the subtype distribution PV:SST:CR among EGFP-positive cells in layer V did not differ significantly from that of the entire cell population in layer V. Taken together, we conclude that cis-element within a *hGPR56 e1m* region contributes to promote GPR56 expression in broad subtypes of GABAergic neurons.

**Figure 5.**
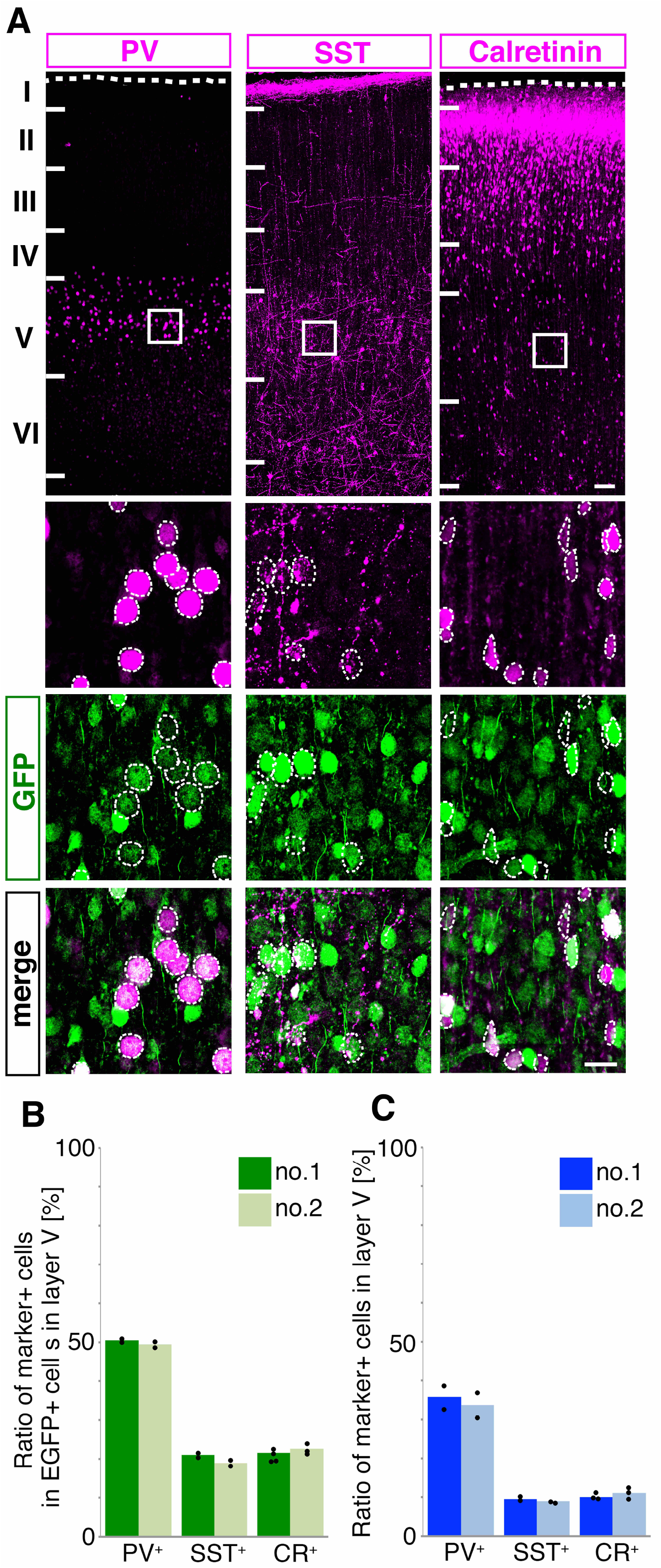
Subtype marker expression in the *hGPR56e1m*-EGFP positive GABAergic interneurons. **(A)** Distribution of *hGPR56e1m*-EGFP and PV, SST or Calretinin in the cortical plate of the transgenic marmoset brain (E113). Scale bar = 1 mm. Enlarged images of the area marked by rectangles in top panels are shown in the lower panels. The dashed ellipses indicate the positive cells of each subtype marker of GABAergic neuron. Scale bars = 50 μm. **(B)** Ratio of PV^+^, CR^+^, or STT^+^ cells among the *hGpr e1m*-driven EGFP^+^ cells in the layer V of transgenic marmoset no.1 and no.2. **(C)** Ratio of PV^+^, CR^+^, or STT^+^ cells among the cells in the layer V of transgenic marmoset no.1 and no.2.

## DISCUSSION

In this study, we examined the role of human *GPR56* e1m minimum promoter by using marmoset as a nonhuman primate model. We developed transgenic marmosets in which EGFP is expressed under the control of human *GPR56* e1m minimum promoter, and examined the profile of EGFP positive cells, especially within the cerebral cortex. In developing marmoset brain, *0.3k hGPR56 e1m*-EGFP showed restricted expression pattern within the endogenous GPR56 expressing regions. While endogenous GPR56 expressed in immature cells including neural stem cells and progenitors, *0.3k hGPR56 e1m*-EGFP expression was rarely detected in such immature cells but detected mainly in developing neurons. These results are consistent with the expression patterns of endogenous mouse Gpr56 and *0.3k hGPR56 e1m*-driven reporter gene in the transgenic mice^7^. We further showed that EGFP in layer V was preferentially expressed in GABAergic neurons. It is well known that, in the developing brain, the anti-CTIP2 antibody preferentially labels a relatively narrow population of cells in layer V of the cerebral cortex. Our present study showed that EGFP^+^ cells in layer V contained a higher percentage of GABAergic neurons and a lesser percentage of CTIP2^+^ cells than pan-GPR56^+^ cells, where the summed percentage of CTIP2^+^ cells and GABA^+^ cells was similar (about 90%) to the EGFP^+^ cells in layer V (Fig. 4E and F). Our data indicates that, even if all of the GABA^−^ EGFP^+^ cells in layer V were glutamatergic neurons, EGFP^+^ cells should contain a higher percentage of GABAergic neurons than that of glutamatergic neurons in layer V (Fig. 4E). GPR56 is classified into adhesion GPRs^29^ that are involved in cell proliferation and migration in the developing brain. Among the genes that mainly contribute to the cell migration of neocortex, homeobox transcription factor DLX families and ARX, which is regulated under DLX, are also involved in differentiation of GABAergic inhibitory neurons at embryonic stages^30,31^. Similar to the case for DLX and ARX, our result suggested that GPR56 expression regulated under the control of the e1m minimal promoter may also contribute to migration and development of GABAergic neurons.

We examined the distribution of principle each subtypes of GABAergic interneurons among the *0.3k hGPR56 e1m*-EGFP positive neurons, and showed that the ratio of PV-positive cells was about two times higher than those of SST-or CR-positive cells. These ratios are roughly consistent with those among the GABAergic neurons in marmoset cortical layer V (Fig. 5B and C). Therefore, it appears that e1m cis-element drives GPR56 expression in GABAergic neurons irrespective of subtype. This may indicate that GPR56 driven by e1m cis-element plays a role in the earlier development of GABAergic neuron before its subtype determination^32^. Taken together, our results support the idea that 15bp deletion within e1m cis-element may reduce the expression of GPR56 in GABAergic neurons in human developing brain.

The preferential expression of e1m-driven EGFP in inhibitory interneurons, and in the early developing ganglionic eminence and later early caudate nucleus is most simply explained by the observation that most inhibitory interneurons originally derive from the several GE’s, including the medial, caudal, and lateral ganglionic eminences. Thus, the e1m element may drive EGFP expression in the progenitor cells of interneurons in the GE during development, as well as in the inhibitory interneurons derived from these structures. In fact, studies in mice suggest that disruption of the e1m element causes a prominent loss of expression in developing GE^7^, consistent with some of our observations here. On the other hand, mutation of the e1m element in mice also disrupts transgene expression in lateral cerebral cortical cells as well, which is less well illustrated by our marmoset transgene. The splice structure of GPR56 is quite dynamic between mouse and primates, and some of these changes may be responsible for these species differences; alternatively they may reflect technical differences in the precise elements of the transgenes used. Since the anti pan-GPR56 antibody recognizes all of the GPR56 isoforms, it is expected that all EGFP-positive cells would also be labeled with the panGPR56 antibody. Indeed, nearly 90% of the EGFP positive cells were positive for pan-GPR56 staining. The remaining EGFP positive cells were negative for the pan-GPR56 signal. This could be explained by the difference in sensitivity between anti-GFP and anti-panGR56 antibodies, although some other regulatory elements may be required to fully recapitulate endogenous marmoset *GPR56* gene expression. Until now, only a few genes have been found, in which mutations in non-cording regions cause epilepsy. For example, the product of *SCN1A* gene is voltage-gated sodium channel Nav1.1 that expresses in GABAergic neuron^33,34,35^. Mutations in its promoter region are reported to reduce *SCN1A* transcription, which causes SCN1A haploinsufficiency. Accordingly, the reduced sodium currents in GABAergic inhibitory neurons may cause hypoexcitability of inhibitory neurons, leading to epilepsy^36^. It is unclear how the deletion within the hGPR56 e1m non-coding region led to epilepsy. An attractive hypothesis would be that, in the case of GPR56, the 15-bp deletion in the cis-regulatory element upstream of the non-coding exon 1m produces intact GPR56 protein^7^, but leads to inaccurate temporal and/or spatial expression of GPR56 in GABAergic neurons along with any potential effects on the development of glutamatergic neurons. Dysfunction of GABAergic neuron development is frequently associated with epilepsy such as with mutations in *DLX* and *ARX*^30,31^. In human, exon 1m of *GPR56* gene is highly expressed in fetal brain compared to the adult brain^7^. Therefore, pathogenic mechanisms of epilepsy associated with patients with a 15-bp deletion within e1m region may be explained in part by the developmental abnormality or dysfunction of GABAergic neurons. Indeed, vigabatrin, a GABA-transaminase inhibitor that is used as a antiepileptic drug, has been reported to relieve symptoms in patients with mutations in *GPR56* gene^12^. Further understanding of the function of GPR56 in GABAergic neurons will help to reveal the precise pathogenic mechanism of epilepsy associated with mutations of *GPR56* gene.

## Methods

### Animals

Experimental procedures were approved by the Animal Care and Use Committees of RIKEN (H30-2-214(3)) and CIEA (11028, 14029, and 15020), and were performed in accordance with their guidelines. Adult common marmosets were obtained from marmoset breeding colonies in CIEA and RIKEN for experimental animals.

### Plasmid constructs and lentiviral production

To construct the *0.3k hGPR56 e1m*-EGFP lentivirus vector plasmid, human *GPR56 e1m* promoter (*0.3k hGPR56 e1m*) was obtained from the plasmid pGL3E-*hGPR56 e1m*-LacZ^7^. CMV promoter sequence of the lentiviral backbone vector, pCS-CDF-CG-PRE (RDB04379, RIKEN, Tsukuba, Japan; a gift from Hiroyuki Miyoshi), was replaced with the *0.3k hGPR56 e1m* promoter. Packaging plasmids, pCAG-HIVgp (mRDB04394) and pCMV-VSV-G-RSV-Rev (RDB04393), were also gifts from Hiroyuki Miyoshi. Lentiviral vector was produced following previously described procedures^37^. In particular, we transfected 30 μg of 0.3 k hGPR56 e1m-EGFP plasmid along with 20 μg HIVgp and 20 μg VSV-G-RSV-Rev packaging plasmids into semi-confluent HEK293T cells in a T175 flask coated with poly-ornitine, using GeneJuice Transfection Reagent (Merck Millipore) according to the manufacturer’s instructions. Six to twelve hours after the transient transfection and the culture at 37° C in a 5% CO_2_ incubator, the medium was replaced with 30 ml FreeStyle 293 Expression Medium (Thermo Fisher Scientific). After 3 days, the culture supernatant containing viral particles was collected, filtered through a membrane with a 0.22 μ m pore size (EMD Millipore, Darmstadt, Germany), and concentrated by ultracentrifugation at 25,000g for 2 hours at 4°C. The viral pellet was then resuspended in 10 μl of ISM1.

### *In vitro* fertilization, early embryo collection, and transplantation

*In vitro* fertilization (IVF) was performed as previously described^21,22^. Donor females’ ovaries were stimulated by intramuscularly injected with human follicle-stimulating hormone (rhFSH, 25IU; FOLYRMON-P injection, Fuji Pharma Co, Tokyo, Japan) for nine days and human chorionic gonadotropin (hCG, 75IU; Gonatropin, ASKA Pharmaceutical Co, Tokyo, Japan) intramuscular injection on day ten then oocytes were collected via follicular aspiration. Collected oocytes were incubated for 24 h at 38°C, 5% CO_2_, 90% N_2_ for in vitro maturation. After incubation, only matured oocytes (metaphase II) were collected and used for IVF. Ejaculated semen was collected non-invasively as described previously^38^. One-cell stage fertilized embryos with two pronuclei were placed in 0.25 M sucrose supplemented PB1 medium (LSI Chemical Medience Corporation, Tokyo, Japan) and the viruses were injected into the perivitelline space using an Eppendorf FemtoJet Express and a Narishige micromanipulator. After cultured beyond 4-cell stage, embryos were transferred to recipient females that had been paired with vasectomized males. After embryo transfer, the recipients were monitored for pregnancy by measuring their plasma progesterone until the pregnancies could be monitored by ultrasound through an abdominal wall.

### Caesarean section

To collect transgenic fertilized embryos from transgenic female I651TgF, subjected to embryo transfer, were paired with intact males for natural embryo collection by nonsurgical uterine flushing. To obtain transgenic embryos at stages E95, 113 and 126, Caesarian Sections were performed in a similar manner as previously described^39^. The pregnant mothers were pre-anesthetized with 0.04 mg/kg medetomidine (Domitor; Nippon Zenyaku Kogyo, Fukushima, Japan), 0.40 mg/kg midazolam (Dormicam; Astellas Pharma Inc, Tokyo, Japan) and 0.40 mg/kg butorphanol (Vetorphale; Meiji Seika Pharma Co, Tokyo, Japan). For maintenance anesthesia during the operation, animals were inhaled 1-3% isoflurane (Forane; Abbott Japan, Tokyo, Japan) via a ventilation mask. After the operation, 0.20 mg/kg antisedan (Atipamezole; Nippon Zenyaku Kogyo, Fukushima, Japan) was injected as an *α*_2_ adrenergic receptor antagonist. On the other hand, to obtain wild type embryos, caesarian sections were performed as previously described^26^. The pregnant mothers were intramuscularly injected with 10 μg/head of atropine sulfate (0.5 mg/ml; Mitsubishi Tanabe Pharma Corporation, Osaka, Japan) followed by with 10 mg/kg of ketamine hydrochloride (Daiichi Sankyo, Tokyo, Japan). 1-3% isoflurane was used for maintenance anesthesia. The embryo and placenta were removed from the uterine by midline laparotomy and then the uterus, abdominal muscles, and skin were sutured. Embryos were anesthetized on ice deeply, dissected in PBS, and the whole brain was removed from the skull.

### Genomic PCR

Genomic DNA was extracted from tissues of wild type (negative control), CMV-EGFP transgenic (positive control) and *0.3k hGPR56 e1m*–EGFP transgenic marmosets using AllPrep DNA/RNA Micro Kit (QIAGEN, Hilden, Germany). For sperm genomic PCR, the CMV-EGFP plasmid (pEGFP; Clontech, CA, USA) was used as a positive control, and the genomic DNA extracted from marmoset ES cells was used as a negative control. They were subject to PCR for transgene detection using the EGFP5-4 (5’-CAAGGACGACGGCAACTACAAGACC-3’) and EGFP3-3es (5’-GCTCGTCCATGCCGAGAGTGA-3’) primers. To detect ß-actin gene, nested PCR was carried out using the first primer set ß-actin 003 (5’-TGGACTTCGAGCAGGAGAT-3’) and ß-actin 006R (5’-CCTGCTTGCTGATCCACATG-3’). PCR was performed for 35 cycles of denaturation at 98 °C for 10 sec, annealing at 65 °C for 10 sec, and elongation at 72 °C for 30 sec.

### RT–PCR

To detect the transgene expression, total RNA was prepared from each tissue and was reverse-transcribed by the SuperScript III First-Ftrand Synthesis System (Thermo Fisher Scientific). PCR was performed using the EGFP5-4 (5’-CAAGGACGACGGCAACTACAAGACC-3’) and EGFP3-3es (5’-GCTCGTCCATGCCGAGAGTGA-3’) primers to detect EGFP gene expression in the tissues, as previous described^21^. To detect ß-actin expression for internal transcript control, the nested PCR was carried out using the first primer set ß-actin 003 (5’-TGGACTTCGAGCAGGAGAT-3’) and ß-actin 006R (5’-CCTGCTTGCTGATCCACATG-3’). All PCR was performed for 35 cycles of denaturation at 98°C for 10 sec, annealing at 65°C for 10 sec, and elongation at 72°C for 30 sec.

### Karyogram analysis

Fluorescent in situ hybridization (FISH) was performed as previously reported (Sasaki et al., 2009) (Chromosome Science Labo Inc, Sapporo, Japan). Peripheral blood samples were obtained from each founder animals. DNA fragment corresponding to a part of the *0.3k hGPR56 e1m*-EGFP was used to produce Cy3-dUTP-labelled probe by the Nick translation method. The common signals among many cells were determined as the insertion sites of *hGPR56 e1m*-EGFP DNA.

### *in situ* hybridization

To generate hybridization probe, about 500 bp cDNA fragment corresponding to 3’ non-coding region of marmoset *GPR56* mRNA was PCR amplified using a cDNA pool derived from marmoset embryonic brain as a template and a set of primers, 5’-ATTCCAATGCTATTTTGCGGGACGTG-3’ and 5’-CAGTTTGTTAGGCAATAACAACAG-3’. Single-color chemiluminescence in situ hybridization (ISH) was performed as previously described^40^. The embryonic brains were drop-fixed in 4% PFA in 100 mM sodium phosphate buffer (PB) pH 7.4 at 4°C more than 24 hours, then replaced into 30% sucrose in 4% PFA in PB. Sections at 40 μm thickness by microtome (Leica, Wetzlar, Germany) were mounted on glass slides and fixed in 4% PFA in PBS for 15 min at room temperature (RT), and treated with proteinase K (Roche, Basel, Switzerland) for 30 min at 37°C. Sections were then fixed again and hybridized with digoxigenin (DIG)-labeled probes (Roche, Basel, Switzerland) at 72°C overnight in hybridization solution (50% formamide, 5xSSC, 1% SDS, 500μg/ml yeast tRNA, 200 μg/ml acetylated BSA, and 50 μg/ml heparin). After washing out excess probe, sections were blocked with 10% lamb serum in TBST for 1 hour at RT and incubated with alkaline phosphatase conjugated to DIG antibody (Roche, Basel, Switzerland) in TBST. Color was developed with a combination of 3.5 mg/ml chromagens nitroblue tetrazolium (Nacalai, Kyoto, Japan) and 1.75 mg/ml 5-bromo-4-chloro-3-indolylphosphate (Nacalai, Kyoto, Japan) in NTMT (100 mM NaCl, 100 mM Tris-HCl (pH 9.5), 50 mM MgCl_2_, 1% Tween 20). Images were taken with an Olympus VS-100 virtual slide system with a 10x objective lens or with a KEYENCE digital microscope, BZ-X700, with a 4x objective lens.

### Transfection and immunocytochemistry

To clone the cDNA of marmoset *GPR56*, marmoset cDNA library was generated. Total RNA was prepared from marmoset brain and was reverse-transcribed by the SuperScript III First-Ftrand Synthesis System (Thermo Fisher Scientific). Marmoset *GPR56* cDNA was amplified by PCR using the primer set marGPR56 5’ end (f)-ATGACTGCCCAGTGCCTCCT and marGPR56 3’ end (r)-GATGCGGCTGGACGAGGTGCT and then re-amplified using the primers marGPR56 *Hin*dIII kozak (f)-CCCAAGCTTGCCACCATGACTGCCCAGTGCCTCCT and marGPR56 *Spe*I 1xHA (r)-GGACTAGTTTAAGCGTAATCTGGAACATCGTATGGGTAGATGCGGCTGGACGAGGTGCT and subcloned into the pCAG-neo. 0.2 *μ*g of the resulting hemagglutinin (HA)-tagged marmoset GPR56 expression vector (pCAG-neo-1xHA-GPR56) was transfected into COS-7 cells cultured in a well of 8 well chamber slide glass using GeneJuice Transfection Reagent (Merck Millipore) according to the manufacturer’s instructions, and the cells were cultured at 37°C in 5% CO2 incubator for 27 hrs. Immunostaining was performed as previously described^41^. Briefly, cultured cells were fixed with 4% PFA-PBS for 20 min at 4°C. The fixed cells were permeabilized with 0.3% Triton X-100-PBS for 15 min at room temperature (RT), incubated with TNB blocking buffer (PerkinElmer) for 1 hr at RT and subsequently incubated with primary antibodies: pan-GPR56 (clone H11) (Millipore MABN310, 1:150) and HA-Tag (3F10) (Merck AB_2314622, 1:1000) overnight at 4°C, followed by incubation with fluorescent-dye-conjugated secondary antibodies: goat secondary antibodies coupled to Alexa 488 (Molecular Probes, 1:200) or Alexa 555 (1:400) for 1.5 hr at RT. Nuclei were counterstained with Hoechst 33258 (10 mg/ml, Sigma-Aldrich). Images were acquired by Carl Zeiss LSM700 confocal microscope using Plan-Apochromat 20x/0.8 M27 objective (Zeiss, 420650-9901), C-Apochromat 63x/1.2W Korr UV-VIS-IR M27 objective (Zeiss, 421787-9970).

### Immunohistochemistry and image acquisition

After dissection, the whole brains were immediately put into 4% PFA in 120mM PB for 24 hours at 4°C, and then were replaced into 30% sucrose in PB at 4°C. After slicing them 50 μm thick with a microtome, immunofluorescence was performed as follows. Antigen retrieval was performed in 0.01 M sodium citrate buffer (pH 6.0) supplemented with 10% (vol/vol) glycerol for 30 min at 85°C, and the samples were left for 30 min back to RT Sections were then washed in PBS, permeabilized in 0.3% Triton-X 100 (wt/vol) in PBS for 30 min at RT, and blocked by TN Blocking buffer (TSA Plus Fluorescence System; PerkinElmer, MA, USA). Primary antibodies were incubated for more than 18 hr at 4°C, and the secondary antibodies were incubated overnight at 4°C.

The following primary antibodies were used: rat monoclonal antibodies to CTIP2 (Abcam ab18465 (25B6), 1:200), somatostatin (Millipore MAB354, 1:50); rabbit polyclonal antibodies to GABA (SIGMA A2052, 1:200), calretinin (Abcam ab16694, 1:100), somatostatin (Abcam ab108456, 1:100), NMDAR1 (Abcam ab17345, 1:100); rabbit monoclonal antibody to TTF1/Nkx2.1 (Abcam ab76013 (clone EP1584Y), 1:100); mouse monoclonal antibodies to pan-GPR56 (Millipore MABN310 (clone H11, IgG_1_), 1:150); goat polyclonal antibodies to GFP (Rockland 600-101-215, 1:300), Parvalbumin (Swant, 1:600); chick polyclonal antibody to GFP (Aves GFP-1020, 1:300). Donkey or goat secondary antibodies coupled to Alexa 488, Alexa 555 or Alexa 647 were used (Molecular Probes, 1:400). For antibodies against GFP56 and Parvalbumin, TSA Plus Fluorescence System (PerkinElmer, MA, USA) was used to enhance the signal. All sections were counterstained with Hoechst33258 (Sigma, 1:1000). Sections were mounted in PermaFluor (Thermo Fisher Scientific) and kept at 4°C. The images were acquired by Carl Zeiss LSM700 confocal microscope using a Plan-Apochromat 10X/0.45 M27 objective (Zeiss, 420640-9900), C-Apochromat 63x/1.2W Korr UV-VIS-IR M27 objective (Zeiss, 421787-9970), and *α* Plan-Apochromat 100x/1.46 Oil DIC M27 objective (Zeiss, 440782-9800). We used ZEN 2009 software (version: 6.0.0.303, Carl Zeiss, Oberkochen, Germany) to acquire z-stack images with a z-interval of either 1–6 μm (10x) or 0.5–1.05 μm (63x, 100x). Section images were taken by an automatic tiling scan system. The images were processed by image J (version: 2.0.0-rc-69/1.52p, U.S. National Institute of Health) and Photoshop CS6 (13.0.6×64, Adobe Inc., CA, USA).

### Determination of layers in the cerebral cortex and cell counting

Each zone in cerebral cortex was defined by the density and direction of nuclei stained with Hoechst 33258 as previously reported^26^. Moreover, in the cortical plate, each layer was defined by the stained patterns of Hoechst 33258 and pan-GPR56 (clone H11). Layer I was identified as a cell-sparse layer. Layer II was identified by radically aligned nuclear packing. The nuclei in layer III also exhibited radial morphology but sparsely rather than in layer II. Layer IV was defined as a densely packed cell layer. Layer V was defined as a relatively cell-sparse layer between the layer IV and VI, and by the expression pattern of CTIP2, which is known to express at high levels in neocortical neurons of layer V^27^. VI was identified as a cell-dense layer located beneath the V. For the identification and quantification of the labeled cells, cell counts were done manually. We first checked 10–13 z-sections within a z-stack of each single channel image and manually marked positive cells on the z-projected image. Then, two z-projected images were merged, and the numbers of overlapping marks were counted as double positive cells. Cell counting was confined to the dorsolateral telencephalon.

## Acknowledgements

We thank Hiroyuki Kanki and Fumiko Ozawa for their contributions to the preparation of probes for *in situ* hybridization. We appreciate Dr. Takuya Shimazaki and Prof. Miho Ohsugi for their advice in the preparation of the manuscript. We also appreciate Prof. Douglass Sipp and Ms. Jaimi Nakajima for proofreading this manuscript. This work was supported by Funding Program for World-leading Innovative R&D on Science and Technology (FIRST) from the Ministry of Education, Culture, Sports, Science and Technology (MEXT), and Brain Mapping by Integrated Neurotechnologies for Disease Studies (Brain/MINDS) from Japan Agency for Medical Research and Development (AMED) (JP19dm0207001, 18dm0207065h0001, 18dm0207002h0005) to T.S., E.S., and H.O. This work was also supported by a grant from Brain science research projects of Center for Novel Science Initiatives of National Institutes of Natural Sciences (NINS) to A.Y.M.

## Author Contributions

A.Y.M. designed the research, conducted the experiments, and wrote the manuscript. K.K., J.O., M.O., and H.M. contributed to the experiments. K.K. and B.I.B. edited the manuscript. T.S. and E.S. conducted experiments. C.A.W., E.S. and H.O. supervised and edited the manuscript.

## Supplementary Information

**Supplementary Figure S1.**
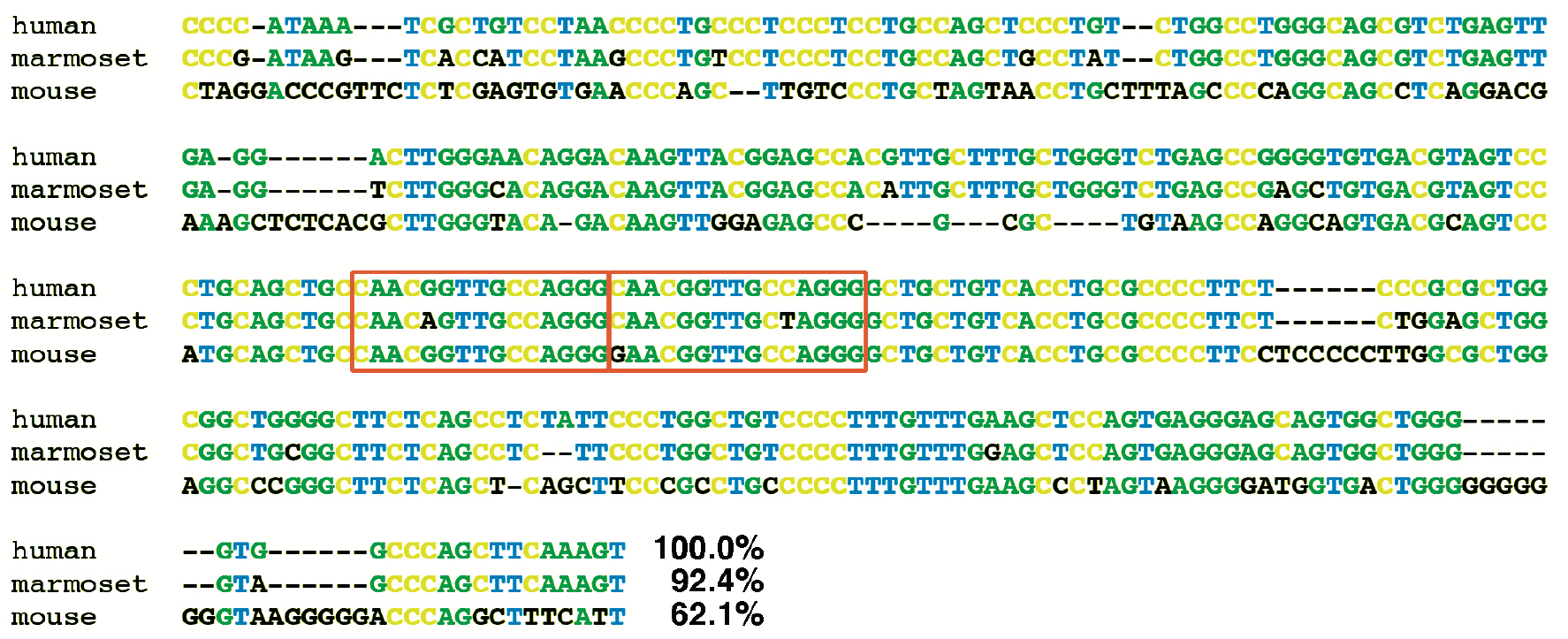
Alignment of 0.3 Kbp e1m sequence of human, marmoset, and mouse. Bases that differ from human sequence are shown in black. The percentages at the end of each sequence indicate the identity to the human sequence. The 15-bp elements are enclosed in orange squares.

**Supplementary Figure S2.**
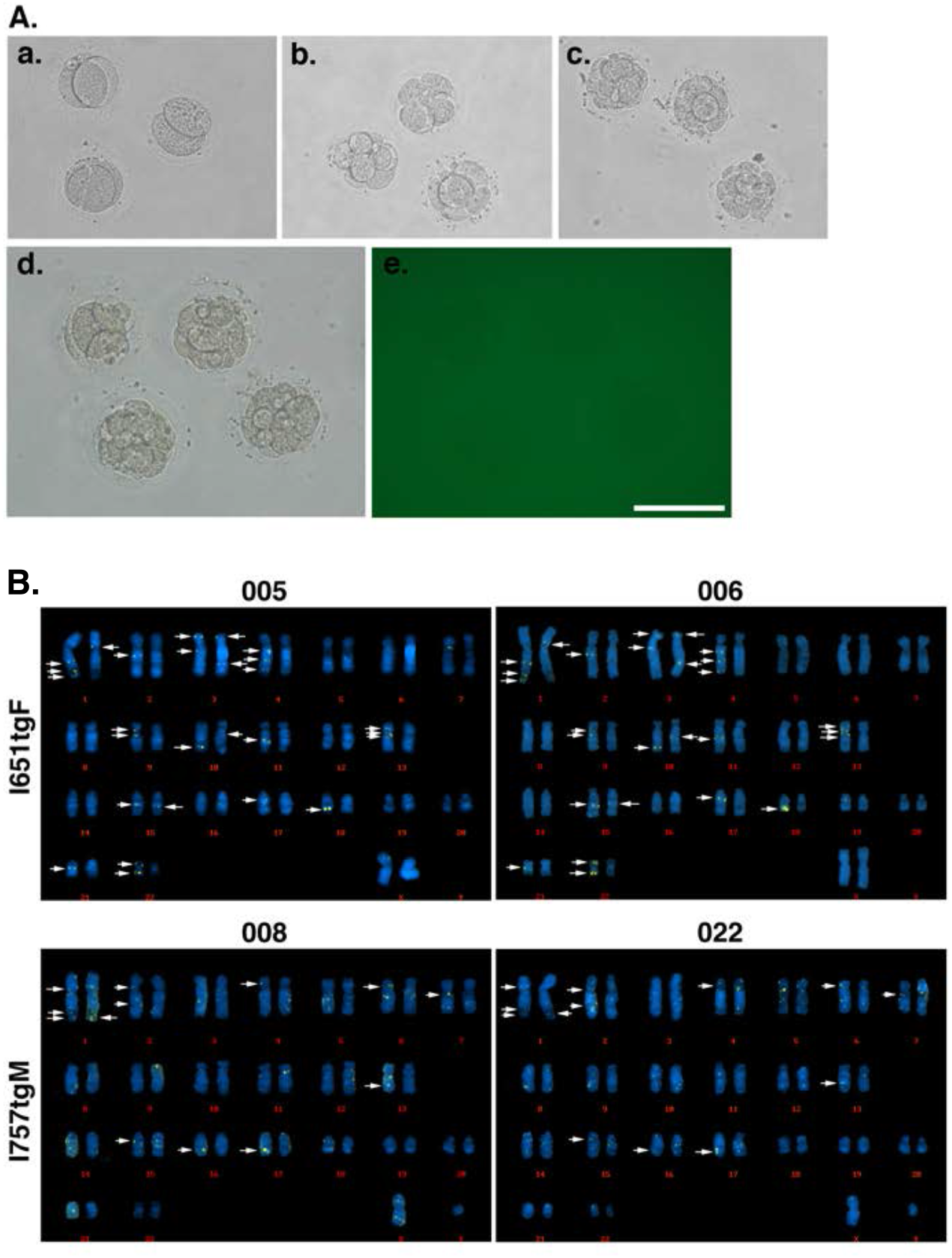
**(A)** Bright field images of marmoset early embryos cultured in vitro for 2 **(a)**, 4 **(b)**, 6 **(c)**, or 7 **(d)** days after lentivector infection. (e) The image of green fluorescence field of (c). *hGPR56 e1m* promoter did not work 7-day cultured early embryos. Scale bar = 100 μ m. **(B)** Genome integration analysis by fluorescence in situ hybridization. The karyograms were prepared from the peripheral blood cells of each founder marmosets, I651TgF and I757TgM. Two sets of the chromosomes from each marmoset are shown as the representatives; #005 and #007 from I651TgF. #0008 and #022 from I757TgM.

**Supplementary Figure S3.**
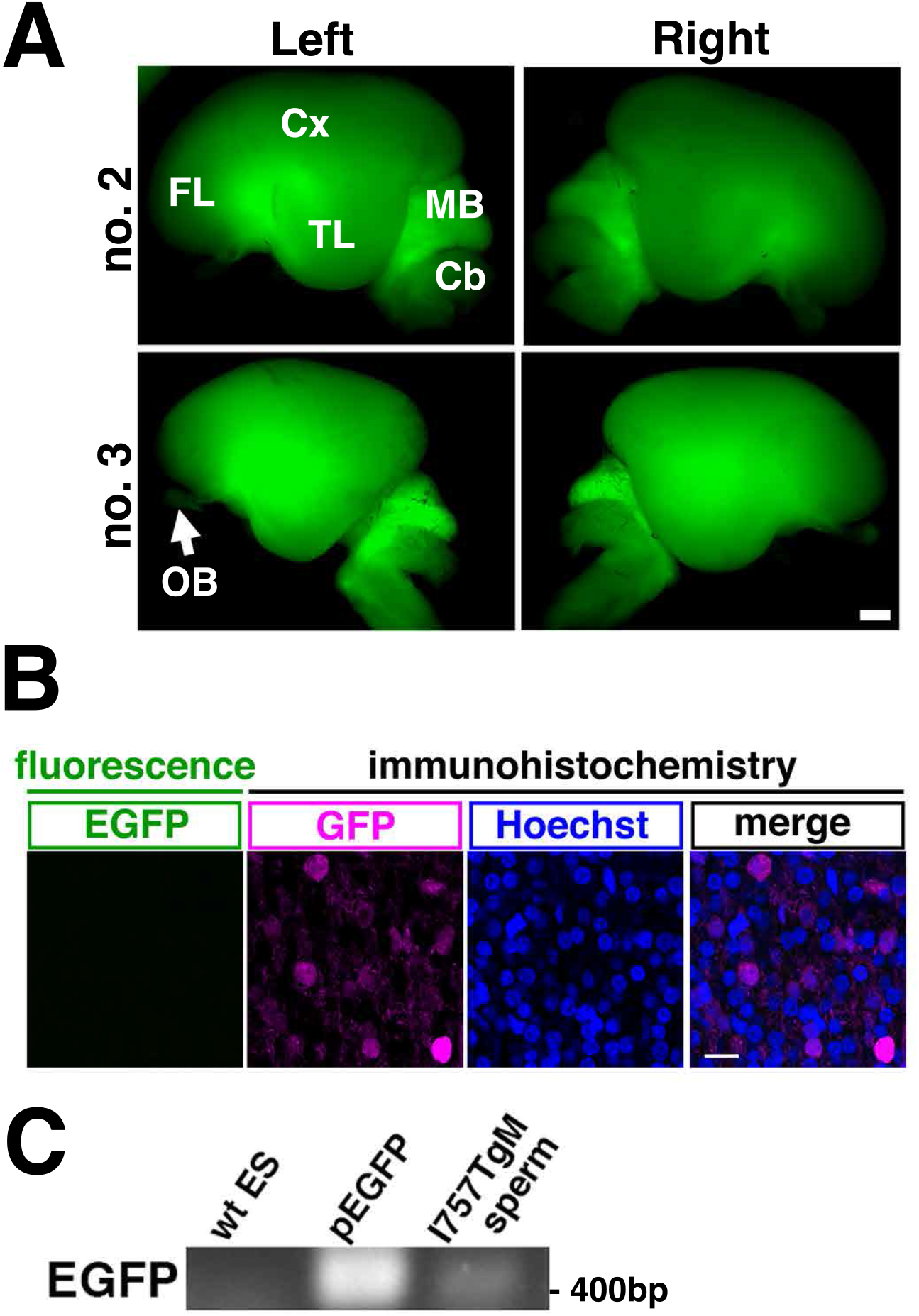
(A) Brains of the E95 fetuses (no.2 and no.3) derived from I651TgF were observed for EGFP fluorescence. Lateral view of the whole brain is shown. Frontal lobe (FL), temporal lobe (TL), cerebral cortex (Cx), midbrain (MB), cerebellum (Cb) and olfactory bulb (OB) are indicated. Scale bar = 1 mm. (B) EGFP fluorescence (green) and immunohistochemistry for GFP (magenta) of the cerebral cortex at E126. DNA was counterstained with Hoechst (blue). Scale bar = 50 μm. (C) Detection of genomic integration of transgene by genomic PCR. Genomic DNA prepared from sperm of I757TgM or wild-type marmoset ES cells (negative control), and pEGFP plasmid (positive control) was used as a template.

**Supplementary Figure S4.**
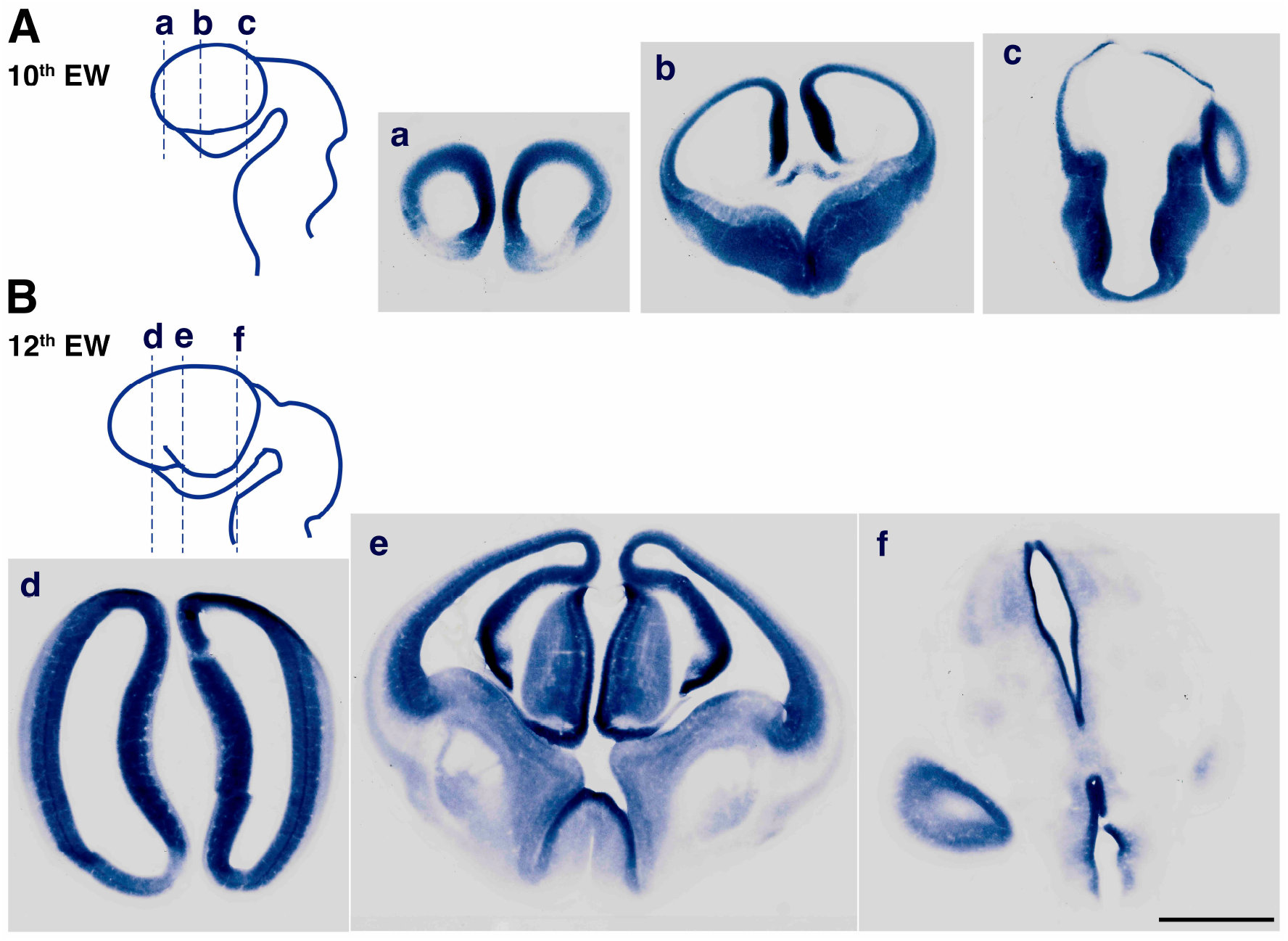
Expression of *GPR56* mRNA in the brain of wild type marmoset embryo. Coronal sections of marmoset brain at the (A) 10^th^ and (B) 12^th^ embryonic week (EW) are hybridized with anti-sense probe for *GPR56* mRNA. Scale bar = 1 mm. Lines on the schema indicate the position of each section.

**Supplementary Figure S5.**
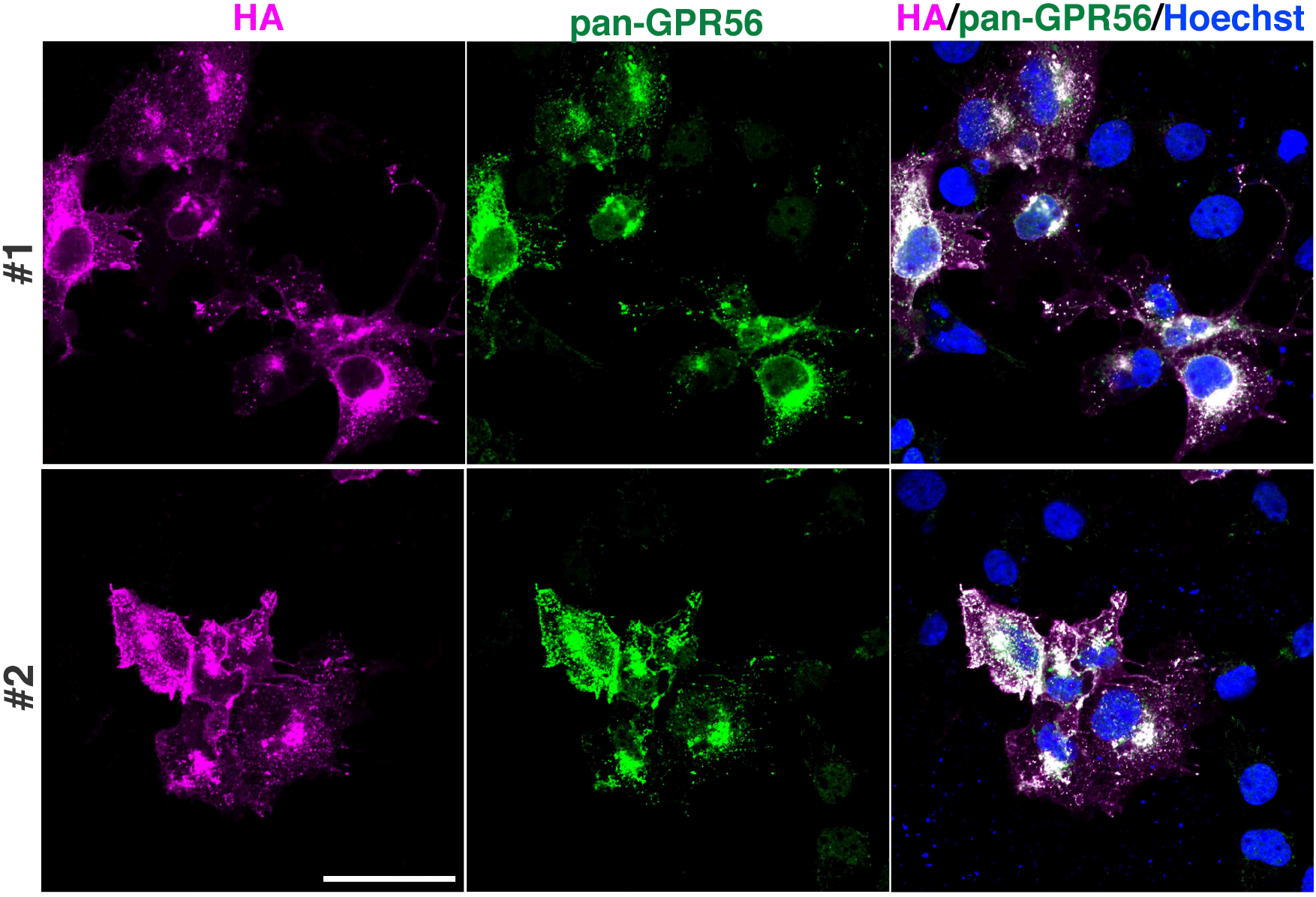
COS cells transfected with HA-tagged marmoset GPR56 expression vector were fixed and stained with anti HA-Tag antibody (magenta), anti pan-GPR56 antibody (green), and Hoechst (blue). Scale bar = 50 *μ*m.

**Supplementary Figure S6 (related to Figure 4C).**
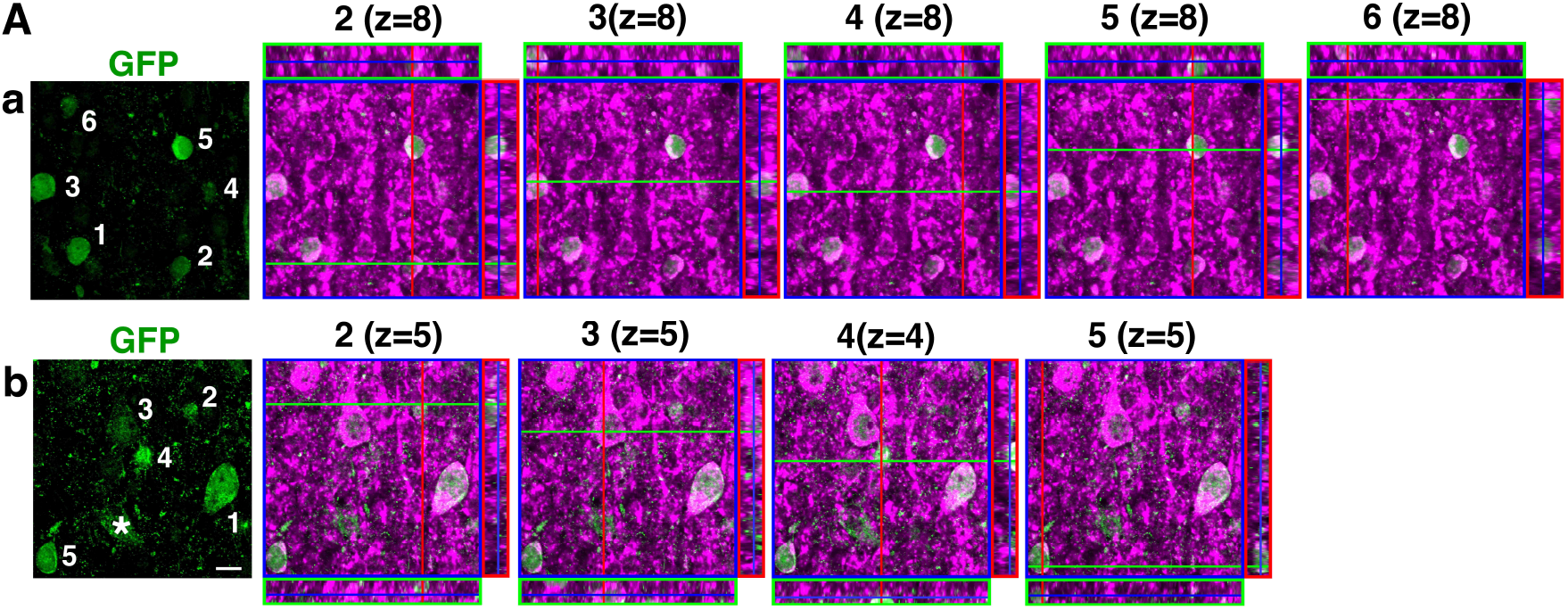
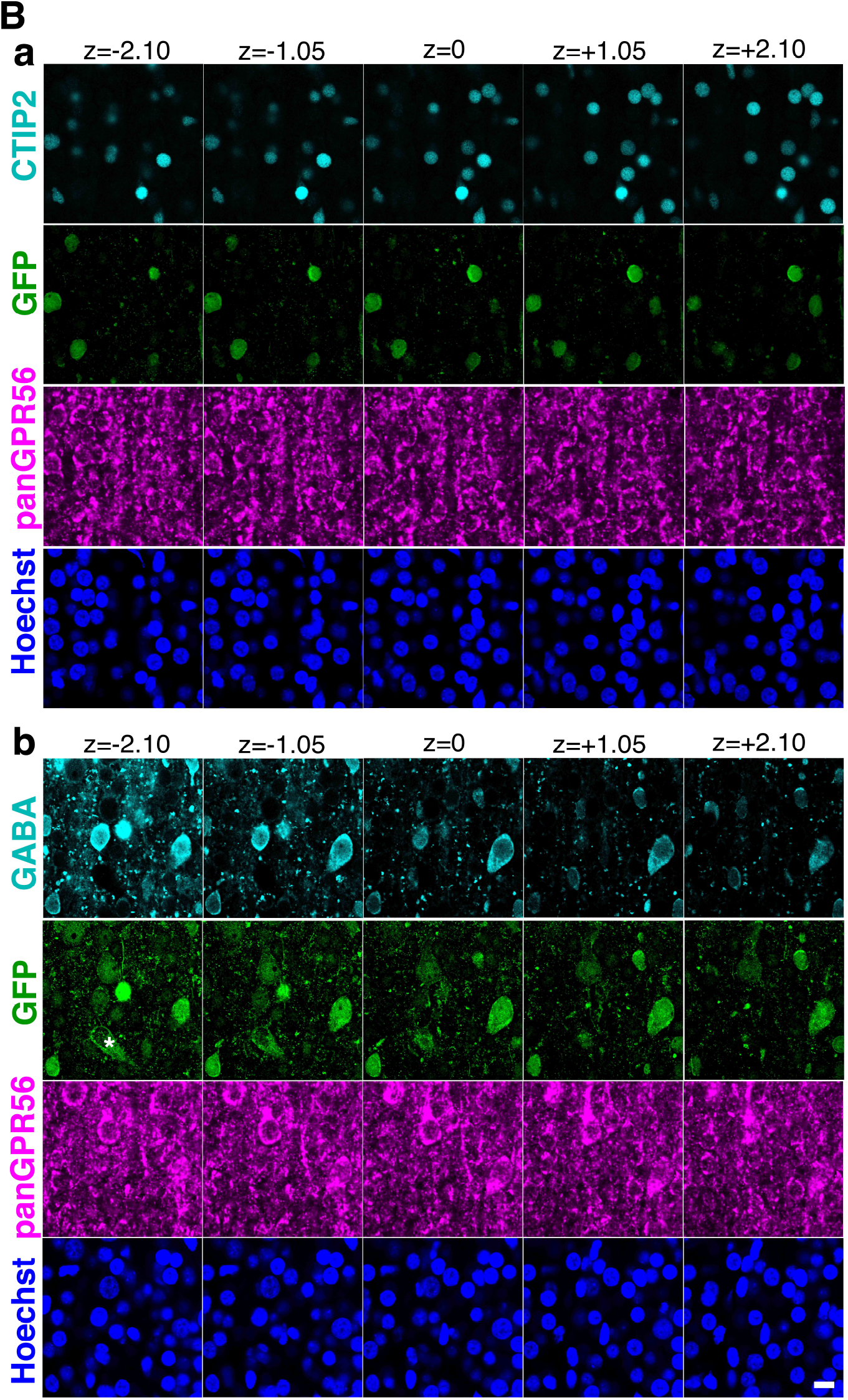
(A) Orthogonal views of *hGPR56 e1m*-EGFP positive (magenta) and pan-GPR56 positive (green) cells in Figure 4C, excluding the one whose orthogonal view image is shown in Figure 4C. Upper (a) and Lower (b) pannels correspond to the Figure 4C a and b, respectively. The left most panels are identical to the GPP panels shown in Figure 4C, but e1m EGFP positive cells are numbered. The cell No. 1 is identical to the cell marked by the arrow in Figure 4C. Z-stack position of each orthogonal view panels is shown in parenthesis. Asterisk indicates blood vessel. (B) Serial z-sections showing the expression of each marker. Orthogonal views and serial z-sections were obtained using ZEN 2009 software (version: 6.0.0.303, Carl Zeiss, Oberkochen, Germany). Scale bars = 10*μ*m.

**Supplementary Figure S7.**
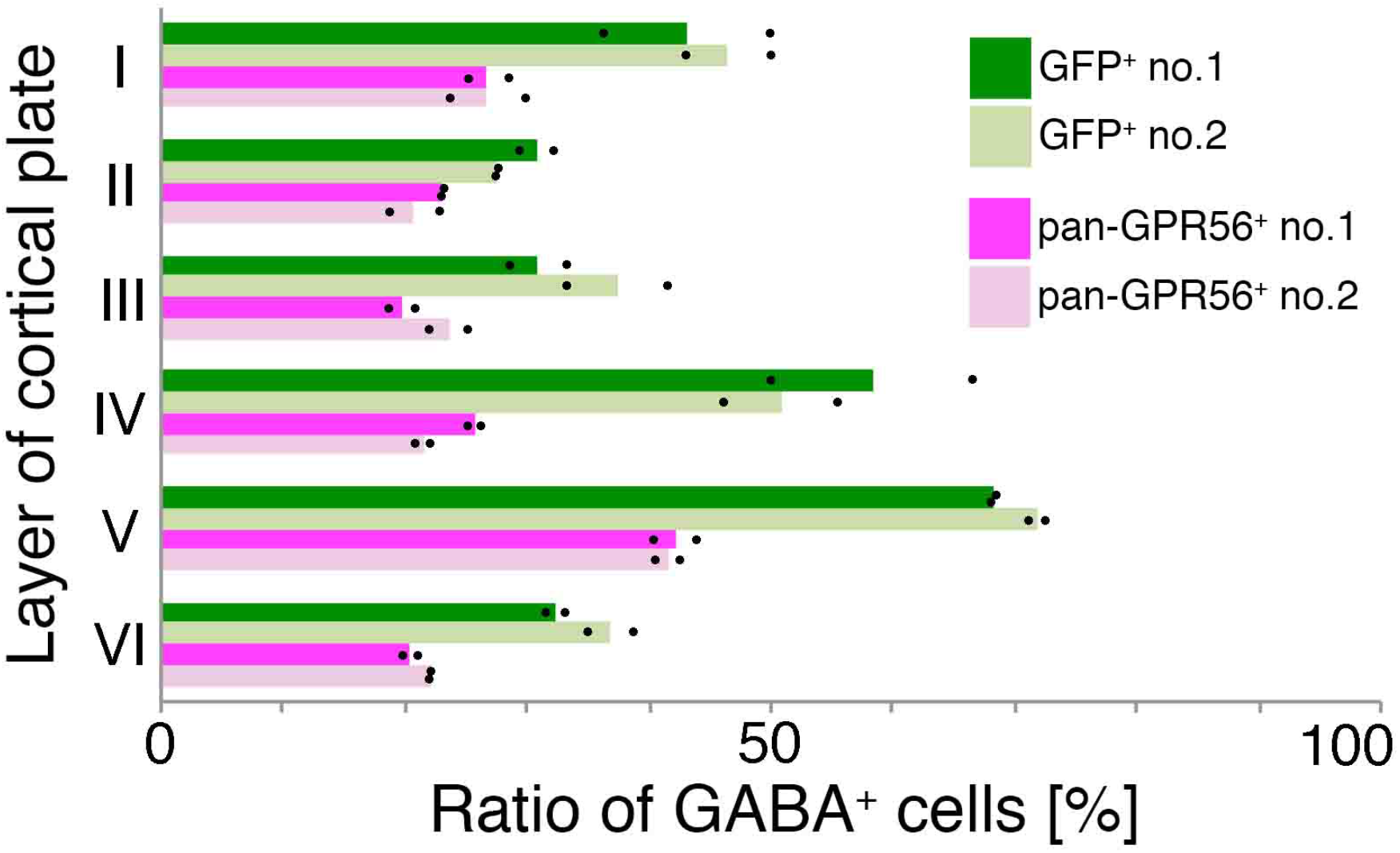
**(A)** Ratio of GABA^+^ cells among the *hGPR56 e1m*-driven EGFP^+^ (green) or pan-GPR56^+^ cells (magenta) in each layer of the cerebral cortex of transgenic marmoset no.1 (dark colors) and no.2 (light colors).

**Supplementary Figure S8.**
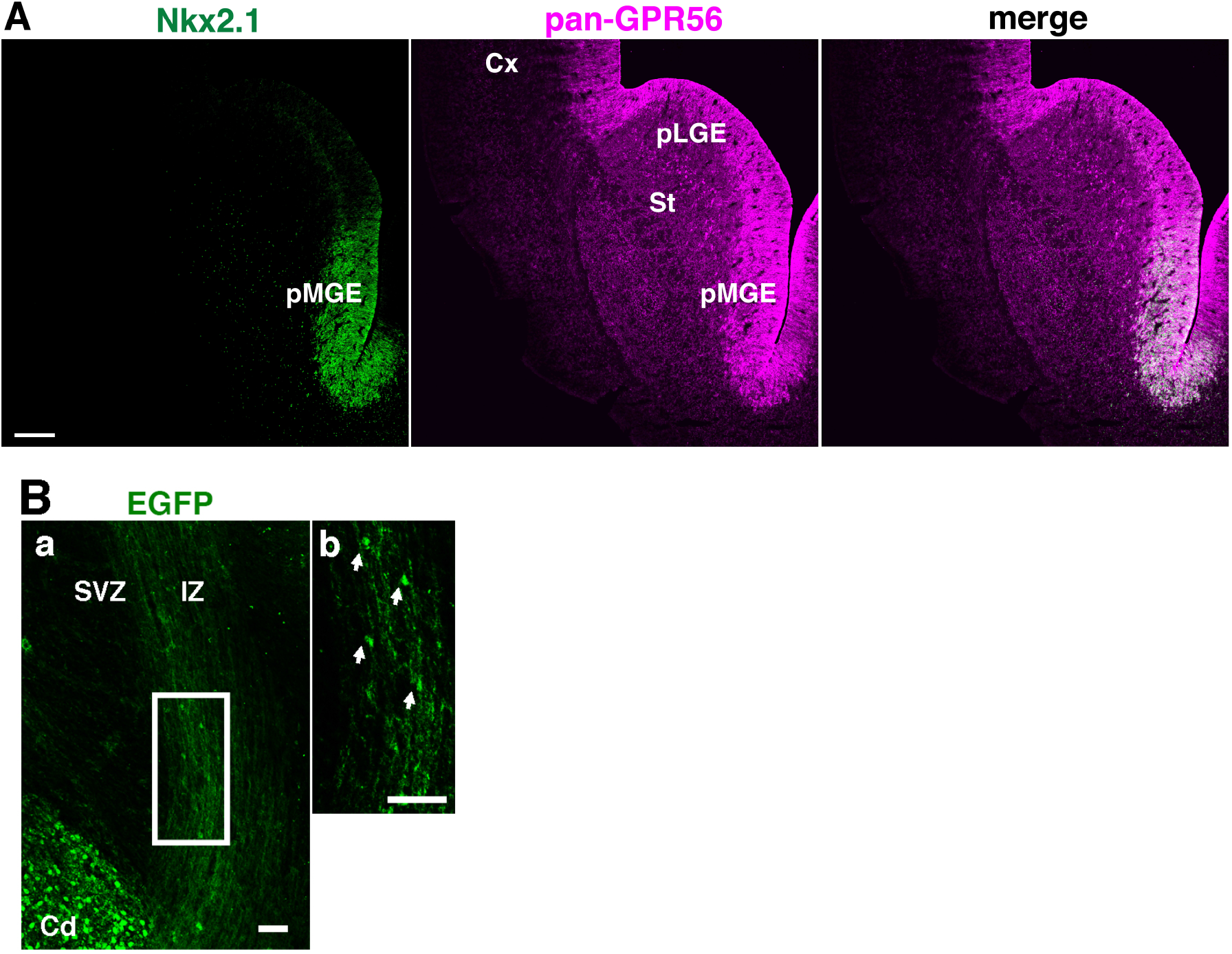
(A) Immunofluorescent staining of coronal section of the marmoset brain at E89 for Nkx2.1 and GPR56. Cerebral cortex (Cx), stratum (St), presumptive lateral ganglionic eminence (pLGE) and presumptive medial ganglionic eminence (pMGE) are indicated. Scale bar = 200*μ*m. (B) Migrating neurons expressing *hGPR56 e1m*-driven EGFP (arrows) in the cerebral cortex of the transgenic marmoset at E95. Subventicular zone (SVZ), intermediate zone (IZ) and caudate (Cd) are indicated. Enlarged image of the area marked by a square in panel a (scale bar=50μm) is shown in panel b (scale bar=100μm).

**Supplementary Figure S9.**
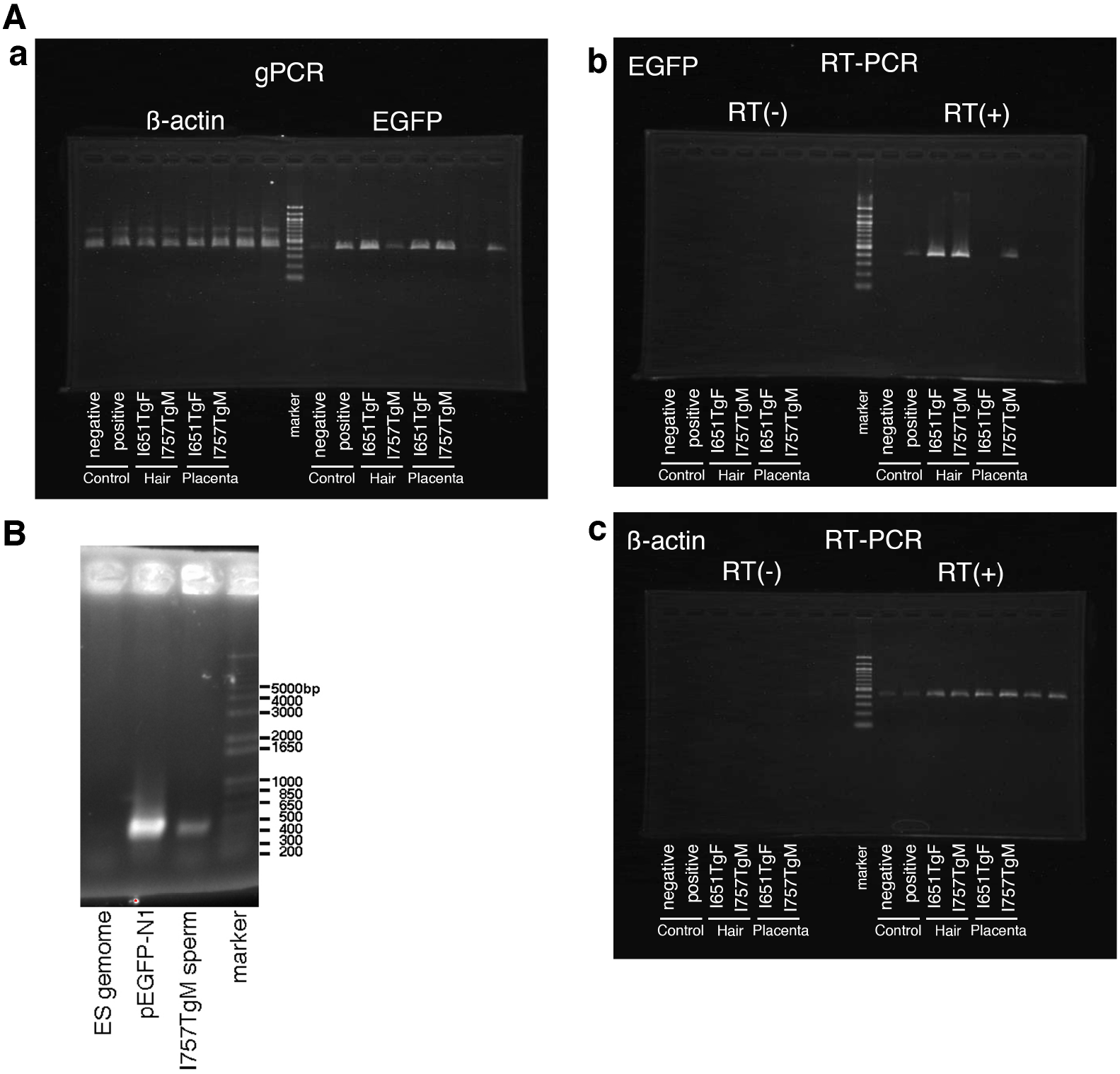
**(A)** Gels scan data of Fig. 1C **(a)** and 1D **(b and c)**. **(B)** Gel scan data of Fig. S3C.

**Supplementary Table S1.** The percentage of *hGPR56 e1m*-EGFP+ cells in each layer. Roman numerals are the number of each layer. The number in the parentheses is the number of sections analyzed.

| individual no. |  | I | II | III | IV | V | VI |
| --- | --- | --- | --- | --- | --- | --- | --- |
| no.1 (12) | mean [%] | 5.3 | 17.7 | 16.3 | 7.4 | 31.0 | 22.3 |
|  | STDEV | 1.4 | 2.2 | 1.5 | 3.1 | 1.0 | 2.7 |
|  | SEM | 0.4 | 0.6 | 0.4 | 0.9 | 0.3 | 0.8 |
| no.2 (13) | mean [%] | 5.7 | 18.5 | 15.7 | 7.0 | 31.5 | 21.5 |
|  | STDEV | 2.0 | 2.1 | 2.8 | 2.3 | 1.0 | 2.2 |
|  | SEM | 0.6 | 0.6 | 0.8 | 0.6 | 0.3 | 0.6 |

**Supplementary Table S2.** The percentage of CTIP2 or GABA immunolabeled neurons among the pan-GPR56 or *hGPR56 e1m*-EGFP expressing neurons in layer V. Dash means no data.

| individual no. |  | pan-GPR56 |  | GFP |  |
| --- | --- | --- | --- | --- | --- |
|  |  | Ctip2 | GABA | Ctip2 | GABA |
| no.1 | % of each sections | 45.8 (71/155) | 40.4 (59/146) | 27.3 (27/99) | 68.36 (67/98) |
|  | (Ctip2+ or GABA+ cells/pan=GPR56+ or GFP+ cells) | 49.7 (79/159) | 37.9 (44/116) | 25.3 (23/91) | 68.0 (51/75) |
|  |  | - |  | 31.3 (10/32) | - |
|  |  | - |  | 30.2 (13/43) | - |
|  | % mean | 47.8 | 39.2 | 28.5 | 68.2 |
| no.2 | % of each sections<br>(Ctip2+ or GABA+ cells/pan-GPR56+ or GFP+ cells) | 52.6 (71/135) | 40.4 (57/141) | 23.5 (20/85) | 71.3 (67/94) |
|  |  | 51.0 (50/98) | 42.6 (63/148) | 25.8 (16/62) | 72.5 (66/91) |
|  |  | 51.0 (51/100) | - | 28.8 (17/59) | - |
|  |  | 47.2 (42/89) | - | 28.3 (17/60) | - |
|  |  | - | - | 30.2 (13/43) | - |
|  |  | - | - | 29.4 (10/34) | - |
|  | % mean | 50.5 | 41.5 | 27.7 | 71.9 |
| total % mean |  | 49.1 | 40.3 | 28.1 | 70.0 |

**Supplementary Table S3.** The percentage of GABA+ cells that coexpress *hGPR56e1m*-EGFP in each layer. Dash means no data.

| layer |  | I | II | III | IV | V | VI |
| --- | --- | --- | --- | --- | --- | --- | --- |
| no.1 | % in each sections (GABA+ cells / EGFP+ cells) | 36.4 (4/11) | 32.3 (10/31) | 33.3 (6/18) | 50(5/10) | 68.4(67/98) | 33.3(10/30) |
|  |  | 50.0 (1/2) | 29.4(5/17) | 28.6(2/7) | 66.7(2/3) | 68.0(51/75) | 31.6(6/19) |
|  | % mean | 43.2 | 30.8 | 31.0 | 58.3 | 68.2 | 32.5 |
| no.2 | % in each sections (GABA+ cells / EGFP+ cells) | 50.0 (3/6) | 27.8 (5/18) | 41.7 (5/12) | 55.6(5/9) | 71.3(67/94) | 35.0(7/20) |
|  |  | 42.8(3/7) | 27.6(8/29) | 33.3 (7/21) | 46.2(6/13) | 72.5(66/91) | 38.9(14/36) |
|  | % mean | 46.4 | 27.7 | 37.5 | 50.9 | 71.9 | 36.9 |
| total % mean |  | 44.8 | 29.3 | 34.2 | 54.6 | 70.1 | 34.7 |
| total counted cell number |  | 26 | 85 | 58 | 35 | 334 | 105 |

**Supplementary Table S4.** The percentage of GABA+ cells that coexpress pan-GPR56 in each layer. Dash means no data.

| layer |  | I | II | III | IV | V | VI |
| --- | --- | --- | --- | --- | --- | --- | --- |
| no.1 | % in each sections (GABA+ cells / pan-GPR56+ cells) | 28.6 (4/14) | 23.3 (7/30) | 18.8(6/32) | 26.3(5/19) | 40.4(59/146) | 19.6(11/56) |
|  |  | 25(2/8) | 23.1 (6/26) | 20.8 (5/24) | 25(2/8) | 37.9(44/116) | 21.2(7/33) |
|  | % mean | 26.8 | 23.2 | 19.8 | 25.7 | 42.2 | 20.4 |
| no.2 | % in each sections (GABA+ cells / pan-GPR56+ cells) | 30.0 (3/10) | 18.8 (3/16) | 22.2 (6/27) | 22.2(6/27) | 40.4(57/141) | 22.2(8/36) |
|  |  | 23.1(3/13) | 22.9(8/35) | 25.0 (8/32) | 20.7(6/29) | 42.6(63/148) | 22.0(13/59) |
|  | % mean | 26.5 | 20.8 | 23.6 | 21.5 | 41.5 | 22.1 |
| total % mean |  | 26.7 | 22.0 | 21.7 | 23.6 | 41.9 | 21.3 |
| total counted cell number |  | 45 | 91 | 115 | 75 | 551 | 184 |

**Supplementary Table S5.** The percentage of PV, SST and CR immunolabeled neurons among the *hGPR56 e1m*-EGFP+ cells in layer V. Dash means no data.

| individual no. |  | PV+ | SST+ | CR+ |
| --- | --- | --- | --- | --- |
| no.1 | % in each sections<br>(Marker+ cells/EGFP+ cells) | 50.0 (16/32) | 20.4 (10/49) | 19.4 (7/36) |
|  |  | 51.2 (22/43) | 21.6 (11/51) | 19.6 (9/46) |
|  |  | - | - | 23.5 (8/34) |
|  |  | - | - | 22.7 (10/44) |
|  | % mean | 50.6 | 21.0 | 21.3 |
| no.2 | % in each sections<br>(Marker+ cells/EGFP+ cells) | 48.8 (21/43) | 18.5 (12/65) | 23.9 (11/46) |
|  |  | 50.0 (17/34) | 19.7 (14/71) | 22.1 (15/68) |
|  |  | - | - | 21.3 (10/47) |
|  | % mean | 49.4 | 19.1 | 22.4 |
| total % mean |  | 50.0 | 20.0 | 21.9 |
| total counted cell number |  | 152 | 236 | 321 |

**Supplementary Table S6.**
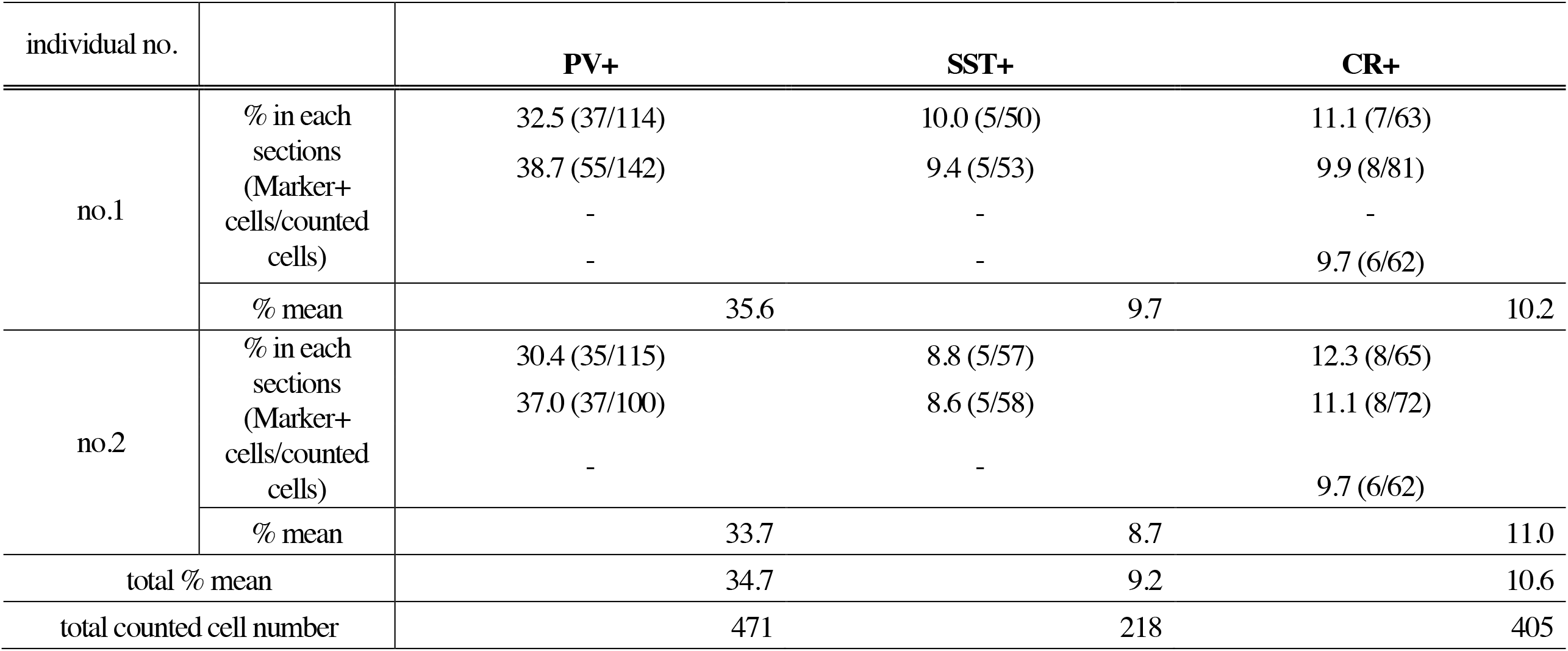
The percentage of PV, SST and CR neurons in layer V. Dash means no data.

| individual no. |  | **PV+** | **SST+** | **CR+** |
| --- | --- | --- | --- | --- |
| no.1 | % in each sections<br>(Marker+ cells/counted cells) | 32.5 (37/114) | 10.0 (5/50) | 11.1 (7/63) |
|  |  | 38.7 (55/142) | 9.4 (5/53) | 9.9 (8/81) |
|  |  | - | - | - |
|  |  | - | - | 9.7 (6/62) |
|  | % mean | 35.6 | 9.7 | 10.2 |
| no.2 | % in each sections<br>(Marker+ cells/counted cells) | 30.4 (35/115) | 8.8 (5/57) | 12.3 (8/65) |
|  |  | 37.0 (37/100) | 8.6 (5/58) | 11.1 (8/72) |
|  |  | - | - | 9.7 (6/62) |
|  | % mean | 33.7 | 8.7 | 11.0 |
| total % mean |  | 34.7 | 9.2 | 10.6 |
| total counted cell number |  | 471 | 218 | 405 |

